# Disinfection exhibits systematic impacts on the drinking water microbiome

**DOI:** 10.1101/828970

**Authors:** Zihan Dai, Maria C. Sevillano-Rivera, Szymon T. Calus, Q. Melina Bautista-de los Santos, A. Murat Eren, Paul W.J.J. van der Wielen, Umer Z. Ijaz, Ameet J. Pinto

## Abstract

Limiting microbial growth during drinking water distribution is achieved either by maintaining a disinfectant residual or through nutrient limitation without the use of a disinfectant. The impact of these contrasting approaches on the drinking water microbiome is not systematically understood. We utilized genome-resolved metagenomics to compare the structure, metabolic traits, and population genomes of drinking water microbiomes across multiple full-scale drinking water systems utilizing these two-distinct microbial growth control strategies. Microbial communities cluster together at the structural- and functional potential-level based on the presence or absence of a disinfectant residual. Disinfectant residual concentrations alone explained 17 and 6.5% of the variance in structure and functional potential of the drinking water microbiome, respectively, despite including samples from multiple drinking water systems with variable source waters and source water communities, treatment strategies, and chemical compositions. The drinking water microbiome is structurally and functionally less diverse and less variable across disinfected systems as compared to non-disinfected systems. While bacteria were the most abundant domain, archaea and eukaryota were more abundant in non-disinfected and disinfected systems, respectively. Community-level differences in functional potential were driven by enrichment of genes associated with carbon and nitrogen fixation in non-disinfected systems and γ-aminobutyrate metabolism in disinfected systems which may be associated with the recycling of amino acids. Metagenome-assembled genome-level analyses for a subset of phylogenetically related microorganisms suggests that disinfection may select for microorganisms capable of using fatty acids, presumably from microbial decay products, via the glyoxylate cycle. Overall, we find that disinfection exhibits systematic and consistent selective pressures on the drinking water microbiome and may select for microorganisms able to utilize microbial decay products originating from disinfection inactivated microorganisms.

## INTRODUCTION

Drinking water systems harbor diverse and complex microbial communities in bulk water, biofilms on pipe wall, suspended solids, and in loose deposits^1–5^. While treatment processes at the drinking water treatment plants (DWTPs) shape the microbial community that leaves the DWTP^6–9^, multiple factors can influence the structure and function of the drinking water microbiome in the drinking water distribution systems (DWDSs). These factors include, but are not limited to, DWDS size, pipe materials and ages, water age within the DWDS and similar factors within premises plumbing (PP) in buildings and homes^10–14^. Managing the microbiological quality of drinking water during transport through the DWDS and into the PP is essential for the provision of safe drinking water. Unwanted microbial growth and/or changes in the drinking water microbiome composition during transit through the DWDS and PP are associated with several detrimental outcomes. For instance, this could lead to proliferation of opportunistic pathogens^15–19^ and an eukaryotic microbes^14, 16, 20, 21^, taste and odor issues^22^, and impact infrastructure via corrosion damage^23, 24^.

Source-to-tap differences in drinking water systems can range from source water type (e.g., surface, ground, reuse water), process configurations at the DWTP, heterogeneity and condition of the DWDS and PP; yet globally there are two fundamental approaches for managing the drinking water microbiome during transport to the consumer^25^. The first and most widely used approach involves maintenance of a disinfectant residual (e.g., chlorine) in the DWDS. This is accomplished by ensuring the water leaving the DWTP has a chlorine residual and/or by using booster stations in large complex DWDSs to compensate for disinfectant residual decay^26^. Disinfectant residuals counteract microbial growth through inactivation, thus ensuring stable microbial concentrations during distribution. While disinfectant residuals are effective in managing microbial growth in the DWDS, there are some key issues associated with them. These include aesthetic and corrosion related problems^25, 27, 28^, but more importantly the formation of harmful disinfection byproducts (DBPs)^29–31^, which are also regulated. Further, there is an increasing recognition that the disinfectant residuals may be associated with selection of some opportunistic pathogens^16, 32^ and antibiotic resistance genes (ARGs) in drinking water^33–35^.

The second approach for managing microbial growth in the DWDS, primarily practiced in parts of western Europe (e.g., Netherlands, Denmark, and Switzerland), involves distribution of drinking water without any disinfectant residuals^36^. These systems focus on minimizing nutrient availability in the DWDS to limit microbial growth using high-quality source waters and/or multi-barrier treatment. While some of these drinking water systems may also use chlorine or other chlorine compounds (e.g., chlorine dioxide) at the DWTP, they ensure that chlorine is not detectable prior to distribution. The efficacy of this approach is supported by evidence that incidences of microbial contamination and associated waterborne illnesses are comparable to systems that maintain a disinfectant residual^25, 37^. This suggest that with appropriate source water quality management, treatment, and well maintained infrastructure, drinking water can be safely distributed without disinfectant residuals^25^.

Despite reports of comparable biological water quality between systems with and without disinfectant residuals, there are a limited number of studies that have systematically compared the microbial community between these two types of systems. Bautista et al (2016)^38^ conducted a meta-analyses study involving collation, curation, and comparison of 16S rRNA gene amplicon sequencing data from previously published datasets. While this study was confounded my methodological differences between datasets being used, the key conclusions were that presence/absence of disinfectant residuals impact microbial community structure and membership and that systems without disinfectant residuals are more diverse than their disinfected counterparts. Recently, Waak et al (2019)^39^ compared biofilms between two drinking water systems, one chloraminated systems and one without a disinfectant residual. Consistent with previous findings they observed higher cell numbers and higher diversity in the system without disinfectant residual, with higher proportional abundance (proportion of total community) of deleterious microbes (i.e., mycobacteria, nitrifiers, corrosion causing bacteria) in the chloraminated system. Both, Bautista et al (2016)^38^ and Waak et al (2019)^39^ utilized gene-targeted assays (i.e., 16S rRNA gene) to probe drinking water microbiome composition and its differences. While gene-targeted assays can provide valuable information on microbial community structure and membership information, they do not provide insight into metabolic differences that may drive the observed differences in community structure. Further, gene-targeted assays can be limited by primer-bias and can result in non-detection of microbial community members. Both challenges can be overcome by utilizing metagenomics which can provide insights into structure and functional potential of a microbial communities without being biases against or towards specific community members. This comes with the limitation that differences between samples/systems emerging from low-abundance microbes may not be detected as this may require ultra-deep sequencing.

We used metagenome analyses and genome-resolved metagenomics to investigate the potential influence of disinfectant residuals on the drinking water microbiomes by comparing drinking water systems from the United Kingdom (with disinfectant residual) and the Netherlands (without disinfectant residual). The goals of this study were (1) to determine the extent to which disinfectant residual shapes the structure and functional potential of the drinking water microbiome, (2) to determine whether the selective pressures of disinfection are conserved across drinking water systems, and (3) identify metabolic pathways underpinning differences in structure and functional potential of the drinking water microbiome. Addressing these questions across different drinking water systems with inherent system-to-system variability (e.g., source water, water chemistry, treatment process, etc.) but one consistent difference - i.e., presence or absence of disinfectant residual - will help highlight disinfection that are conserved and thus generalizable across systems.

## MATERIALS AND METHODS

### Sample collection and processing

Drinking water samples were collected from 12 drinking water systems in Netherlands (n=5) between October to December 2013 (Non-disinfected, i.e. ND) and the United Kingdom (n=7) between April to August in 2015 (Disinfected, i.e. D). Samples were collected at two to four locations in each DWDS which resulted in 23 D and 18 ND samples. A total 15 liters of water was filtered through three sterile Sterivex filters with 0.22*μ*m pore size polyethersulfone membrane (EMD Millipore™ SVGP01050) using a peristaltic pump (Watson-Marlow 323S/D) to harvest microbial cells. Immediately after filtration, the membranes were removed aseptically from the Sterivex cartridge, cut into pieces and then transferred to Lysing Matrix E tubes. The membranes were stored at 4°C for 24 hours or less before being transported to the laboratory and stored at - 80°C. Further details of sample treatments and preservation are described in Sevillano-Rivera et al.^35^, along with detailed description of chemical analyses. Briefly, Orion 5 Star Meter (Thermo Fisher Scientific, Waltham, MA) was used to measure temperature, pH, conductivity and dissolved oxygen, total chlorine, and phosphate was also determine on-site using DR 2800 VIS Spectrophotometer (Hach Lange, the UK) and EPA approved HACH kits. Nitrogen species were measured according to standard method, 4500-NH3-F for ammonia, 4500-NO2-B for nitrite, and 4500-NO3-B for nitrate respectively in laboratory^40^, while total organic carbon (TOC) was determined using Shimadzu TOC-LCPH Analyzer (Shimadzu, Kyoto, Japan).

### DNA extractions

The total genomic DNA was extracted directly from filter membranes using Maxwell16 DNA extraction system (Promega) and LEV DNA kit (AS1290, Promega, Madison, WI, US). The filters with collected biomass in lysing matrix E tubes were incubated with 300*μ*L of lysing buffer and 30*μ*L of Proteinase K and incubated at 56°C. A total of 500*μ*L of chloroform:isoamyl alcohol (24:1, pH 8.0) was added to the tube, vortexed and this was followed by bead beating for 40 s at 6 m/s using a FastPrep 24 instrument (MP Biomedicals, Santa Ana, CA, USA), and centrifugation at 14,000g for 10 min. The bead beating and centrifugation steps were repeated twice more with transfer of supernatant to clean tube followed by replacement of the aqueous phase with fresh lysing buffer. The aqueous phase was then subject to DNA purification using the Maxwell LEV DNA kit. The extracted DNA was quantified using Qubit HS dsDNA assay with Qubit 2.0 Fluorometer (Life Technologies, UK). Negative controls consisting of reagent blanks (no input material) and filter blanks (filter membranes from unused Sterivex filters) were processed identically as the samples for DNA extraction. Genomic DNA extracted from mock community, consisting of 10 organisms, detailed previously^35^, was spiked into negative controls extracted (n=8) from the reagent and filter blanks. These negative controls were also included in following library preparation and high-throughput sequencing (see below).

### Library preparation and Illumina sequencing

Sequencing libraries were prepared using the Nextera XT DNA Sample Preparation Kit (Illumina Inc.). All DNA extracts (including negative controls) were cleaned up with HighPrep PCR magnetic beads (MagBio Inc.) to remove short fragments after library preparation and quantified with qPCR according to Illumina guidelines. All libraries were pooled together in equimolar proportion and pooled library was quantified with Qubit HS dsDNA assay and further concentrated using HighPrep PCR magnetic beads (MagBio Inc). Metagenomic sequencing on prepared libraries were performed on four lanes of Illumina HiSEQ 2500 flow cell (2×250-bp read length, Rapid Run Mode) at University of Liverpool Centre for Genomic Research (Liverpool, UK).

### Metagenomic read based analyses

The FASTQ files were trimmed using Cutadapt v1.2.1 (Martin 2014) with a ‘-O 3’ flag, and Sickle v1.200 (Joshi and Fass 2011) using a threshold of window quality score (≥ 20) and read length after trimming (≥ 10 bp). A further trimming was applied using Trimmomatic v0.35^41^ to remove any remaining Illumina Nextera adaptors and trim reads according to quality score with a 4-base wide sliding window and a minimum average quality score of 20 and singlet reads were excluded in downstream analyses as well. To estimate metagenome diversity and coverage for each sample, Nonpareil 3.0^42^ was used in kmer mode on the quality filtered reads. Diversity and coverage information for each metagenome was estimated using command ‘Nonpareil.set()’ in R package ‘Nonpareil’. MicrobeCensus^43^ was used on quality trimmed reads to estimate average genome size across samples with flag ‘-n 100000000’ for all samples. To eliminate the potential effects of bacteria with small genomes (i.e., *Patescibacteria*) on average genome size estimations, pre-processed reads were mapped against 12 *Patescibacteria* metagenome assembled genomes (MAGs) from this study (see below) and 1,037 *Patescibacteria* genomes from GTDB-tk^44^. The reads mapped in proper pair to *Patescibacteria* were removed using samtools (‘-F2’ flag). MicrobeCensus was used again to estimate average genome size using the same parameters.

### Metagenome assembly and mapping

Filtered pair-ended reads were then pooled from each drinking water system for co-assembly, which resulted in 12 paired-end FASTQ files for co-assembly, including seven from disinfected (Dis) and five from non-disinfected (NonDis) systems. *De novo* co-assembly was performed using MetaSPAdes v3.10.1^45^ with recommended k-values for 2×250bp reads (21,33,55,77,99,127). All scaffolds shorter than 500bp were discarded and UniVec_Core build 10.0 (National Center for Biotechnology Information 2016) was used for contamination vector screening and any scaffold with a significant hit to the UniVec database was removed. Reads from each samples were then mapped back to the filtered scaffolds using BWA-MEM v0.7.12 with default settings^46^.

To eliminate the scaffolds that may have originated from sample or post-processing contamination, reads from negative controls were first mapped back to mock community genomes using BWA- MEM v0.7.12^46^, and all reads not mapped in proper pair were extracted using samtools v1.3.1 (Li et al. 2009) with ‘-f2’ flag and were considered “contaminant reads”. Sample reads (S), contaminant reads (C) and negative control reads (NC) were mapped back to filtered scaffolds in each co-assembly. Properly-paired mapped reads were extracted using samtools v1.3.1 with ‘-f2’ flag from the BAM files. Relative abundance and normalized coverage deviation of each scaffold was calculated using reads from samples and those identified as contaminant reads in negative controls:

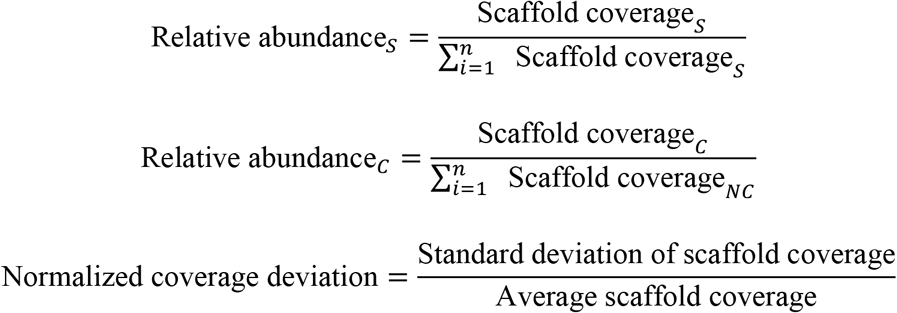

To distinguish true scaffolds from contamination, relative abundance (RA) and normalized coverage deviation (NCD) estimated using sample reads (S) and contaminant reads (C) was compared for all scaffolds:

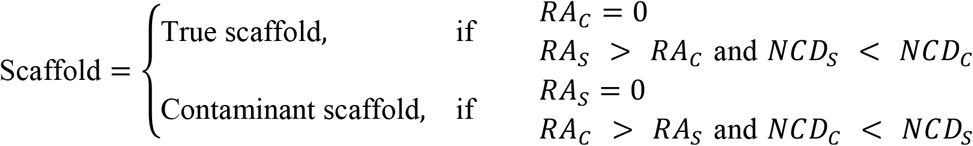

True scaffolds, the scaffolds with higher RA and lower NCD in samples compared to negative controls, were kept for downstream analyses while contaminant scaffolds were excluded from all further analyses.

### Nucleotide and protein composition analyses

MASH v1.1.1^47^ was used to estimate the dissimilarity between samples using quality filtered reads (with ‘-r’ and ‘-m 2’ flags) and dissimilarity between drinking water systems using true scaffolds with the sketch size of 100000. Prodigal v2.6.3^48^ was used to identify open reading frames (ORFs) in the true scaffolds and translate ORFs to protein-coding amino acid sequences. Following prediction and translation, HMMER v3.1b2^49^ was used to annotate ORFs against the Pfam database v31.0^50^ with a maximum e-value of 1*e* − 5 and curated bit score thresholds (the gathering thresholds). Subsequently, MASH distances were calculated between drinking water metagenomes using predicted ORFs, as well as Pfam annotated proteins with the sketch size of 100000 and ‘-a’ flag.

### Taxonomic classification and phylogenetic analyses

The program ‘cmsearch’ was implemented in Infernal v1.1.2^51^ to search scaffolds against SSU rRNA covariance models (CMs) for bacteria, archaea and eukaryota; these are default models used by SSU-ALIGN v0.1^52^ using HMM-only approach and only significant hits were considered. The results were filtered according to length (≥ 100 bp alignment) and e-value (<1*e* − 5). SSU rRNA sequences detected in contaminant scaffolds were removed and if more than one SSU gene sequence was found on a single scaffold, only the longest SSU gene sequence was retained. Relative abundance of each SSU gene sequence was calculated for each sampling location as follows:

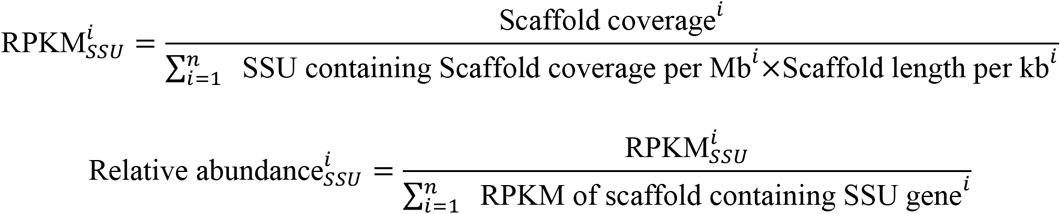

SSU rRNA gene sequences were classified using Mothur v1.33.3 (Schloss et al. 2009) with SILVA database^53^ (Release 132) with a minimum confidence threshold of 80%.

### Annotation and Comparison of functional orthologies and modules between samples

The protein-coding sequences were searched against KOfam, a HMM profile database for KEGG orthology^54^ with predefined score thresholds using KofamScan^55^. Only KEGG orthologies (KO) identified on scaffolds with (> 1x) coverage for each sample and those detected more than once across samples within a single drinking water system were retained for further analyses. Average read count for each KO was calculated using scaffold coverage, average length of reads mapped, and total number of reads mapped to each scaffold in a sample using above equations. To assess functions at KEGG module level, BRITE hierarchy file was retrieved from KEGG website, and KO’s were categorized into KEGG modules. The abundance of KEGG module in each sample was calculated using the median abundance of the detected KEGG orthologies within each module. The completeness of each KEGG module was calculated using ‘KO2MODULEclusters2.py’.

### Metagenome binning and refining

Anvi’o (versions: v2.2.2, v2.4.0, v4 and v5.1)^56^ was used for metagenome binning and refining. Briefly, CONCOCT^57^ integrated in Anvi’o was used to cluster scaffolds (longer than 2500 bp) into metagenome bins using tetra-nucleotide composition and coverage information across all samples within each metagenomic co-assembly. The ‘merge’ method of CheckM v1.0.7^58^ was used to identify the bins that that may emerge from the same microbial population, but may have been separated during automated binning process. Following merging of compatible bins, RefineM v0.0.21^59^ was used to automatically refine bins according to genomic properties (i.e., the mean GC content, tetra-nucleotide signature and coverage) and taxonomic classification. The completeness and redundancy of each refined bin was estimated using CheckM based on collections of lineage specific single-copy genes resulting in a total of 154 bins with greater than >50% completeness. Among these bins, 130 bins had a redundancy of <10% redundancy, while 24 bins are with >10% redundancy. Further manual curation of these bins was performed using Anvi’o, resulting in 156 curated metagenome assembled genomes (MAGs). The 156 MAGs were de-replicated using dRep v2.2.2^60^ and MAGS with >10% redundancy were discarded which resulted in 115 dereplicated MAGs with completeness >50% and reduncancy <10%. All raw sequencing data and dereplicated MAGs are available on NCBI at BioProject number PRJNA533545.

### MAG-level analyses

Taxonomy assignment of MAGs was performed using GTDB-Tk v0.1.3^44^ with the flag ‘classify_wf’. Genome sizes of MAGs were estimated by multiplying the number of nucleotides in the MAG with the inverse of the CheckM estimated completeness. The MAGs were annotated using the HMM profile database for KEGG orthology with predefined score thresholds using KofamScan^55^. The KO’s for each MAG were then categorized into modules based on BRITE hierarchy file retrieved from KEGG^54^, and the completeness of KEGG modules in each genome was calculated using script ‘KO2MODULEclusters2.py’. Anvi’o was used to extract a collection of 48 single-copy ribosomal proteins^61^ from each MAG using ‘anvi-get-sequences-for-hmm-hits’ with a maximum number of missing ribosomal proteins of 40. Subsequently, a phylogenetic tree was reconstructed using concatenated alignment of ribosomal proteins sequences using FastTree v2.1.7^62^. Interactive Tree Of Life (iTOL) v4 (Letunic and Bork 2007) was used to visualize the phylogenetic tree.

Program ‘Union’ in EMBOSS v6.6.0.0^63^ was used to concatenate all scaffolds in each MAG into a single sequence. Reads from all samples were cross-mapped to all MAGs using BWA-MEM v0.7.12 with default settings and proportion of each nucleotides in MAG covered by at least 1x coverage was determined using BEDtools^64^. A MAG was considered detected in a sample if ≥25% of its bases were covered by at least one read from the corresponding sample. This approach was used to determine whether MAGs were detected in all the samples. Further, the MAGs were binned into four categories based on their detection/non-detection within samples. Specifically, MAGs were divided into “D-only” if there were detected in ≥20% of the samples from the disinfected systems and not detected in any samples from the non-disinfected systems, “ND-only” if there were detected in ≥20% of the samples from the non-disinfected systems and not detected in any samples from the disinfected systems, “both” if there were detected in ≥20% of disinfected and non-disinfected systems, while the remaining MAGs were classified in the “other” category. Subsequently, reads from all samples were cross-mapped back to all the MAGs using BBMap v38.24^65^ with a minimum identity of 90%, and ‘ambiguous=best’ and ‘pairedonly=t’ flags. After filtering for detection (see above), reads per kilobase of per million reads (RPKM) for each MAG and each sample were calculated using equation:

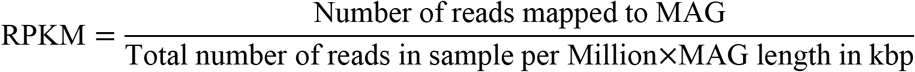

### Statistics

Differences between disinfected and non-disinfected systems for (1) Mash distances distributions, (2) relative abundances were determined using Permutational ANOVA and Pearson’s correlations between pairwise mash distances were estimated in R. BioEnv in “sinkr” (*https://github.com/menugget/sinkr*) and “vegan”^66^ packages were used to identify environmental parameters (i.e., water chemistry) and their combinations that explain the differences in the structure (i.e., Mash distances between samples estimated using reads) and functional potential (i.e., Bray Curtis distance estimated between samples using KO abundance (i.e., RPKM). BioEnv permutes through 2^n-1 possible combination of selected environmental parameters, 511 combinations in this case, and selects the combinations of scaled environmental variables which capture maximum correlation between dissimilarities of community datasets water chemistry and microbial community structure or functional potential. While, BioEnv analyses identified combination of variables that are highly correlated with differences in microbial community structure of functional potential, it does not identify the proportion of variance in microbial community structure of functional potential explained by individual variables or their combination. To this end, we used distance-based redundancy analysis (dbRDA) to perform constrained ordinations on community structure and functional potential to bypass the limitation of usual RDA and CCA, which can only use Euclidean distance measure. Function dbrda() from ‘vegan’ package was used with pairwise Mash distances calculated between samples estimated using reads based Mash distance and Bray-Curtis distances based on KO RPKM to investigate relationships between the environmental variables and community data on both nucleotide composition and KO level. The function varpart() in the vegan package was used to determine the fraction of variation captured parameters identified as significantly associated with read-based Mash and KO relative abundance-based Bray-Curtis distance matrices. DeSeq2 package v1.18.1^67^ was used to identify differentially abundant KEGG modules between disinfected and non-disinfected systems by only considering KEGG modules with a maximum of one block missing and equal to or greater than 50% complete. The median scaffold-length normalized read count of KO’s within each module were used in DESeq2 analyses with a maximum adjusted *P*-value of 0.005.

## RESULTS AND DISCUSSION

### Water quality parameters across disinfected and non-disinfected DWDS

Sampling was conducted in seven DWDSs with disinfectant residual between April-August of 2013 and at five DWDSs without disinfectant residual between October-December 2015. The water chemistry varied between the DWDSs considering they were supplied by different DWTPs, our sampling campaign also captures seasonal differences between locations (Figure 1) (Table S1). Specifically, water temperatures were higher (~5°C) for the disinfected samples compared to the non-disinfected samples. While the pH, DO, nitrogen species (i.e., ammonium and nitrate) and TOC measurements were not significantly different between disinfected and non-disinfected samples, the measured phosphate and total chlorine concentrations were significantly different (p<0.05). Specifically, the average total chlorine concentrations in disinfected systems 0.37 mg Cl_2_/l (range: 0.1-0.73 mg Cl_2_/l) while no disinfectants residuals were measurable in the non-disinfected systems. The average phosphate concentrations were 2.3 mg PO_4_^3−^/l while no phosphate was measurable in non-disinfected samples. Phosphate was higher in the disinfected systems as it is likely to be used for corrosion control^68^. While we were unable to obtain information on source water type (i.e., ground vs surface water) used for production of drinking water supplied to the sampled DWDS, conductivity measurements suggested DWDS in both systems were supplied by a DWTPs drawing from surface and ground water sources (Figure 1).

**Figure 1:**
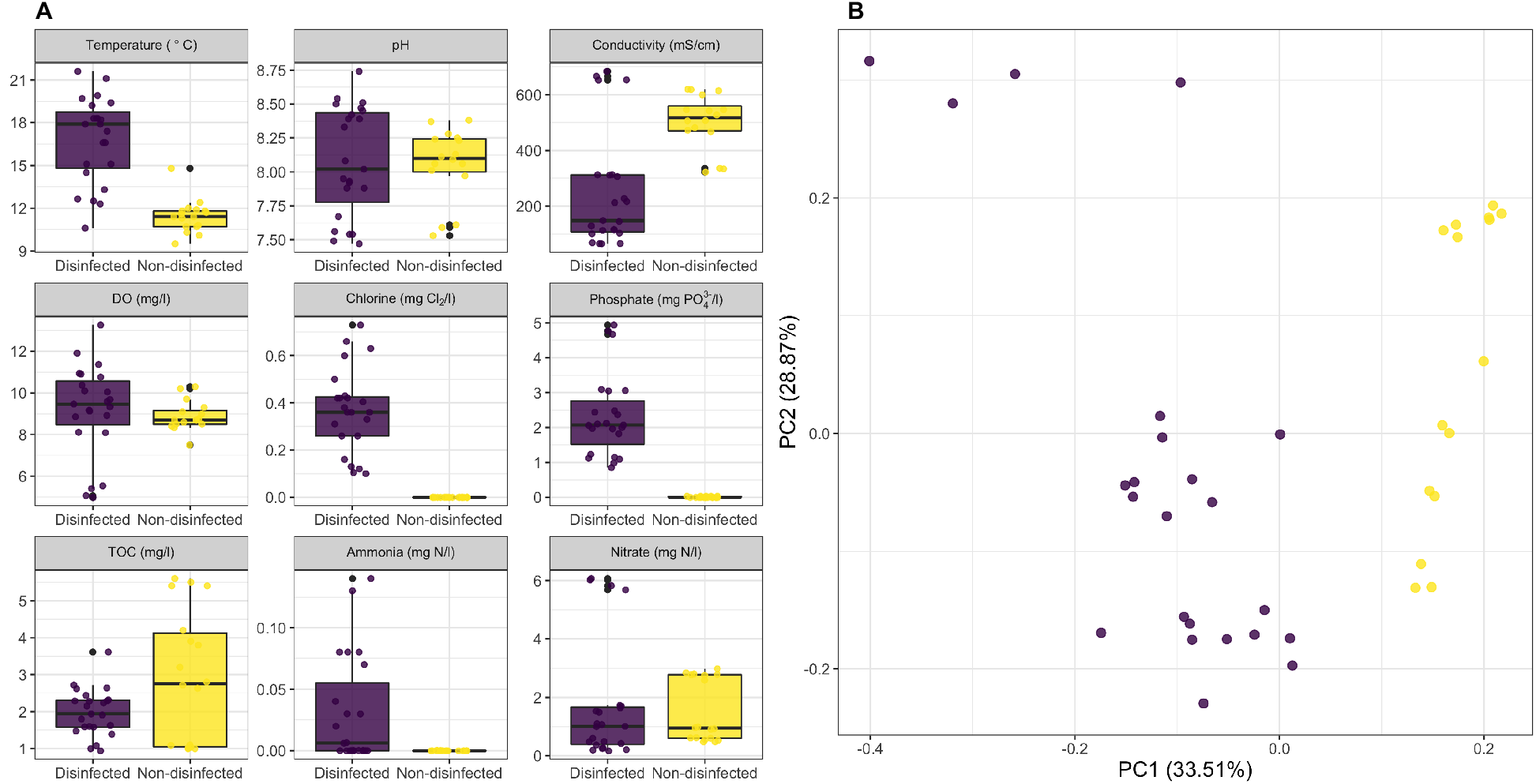
Summary of water chemistry parameters measured for samples collected from disinfected (purple) and non-disinfected systems (yellow). (B) Principle component analyses using Euclidean distances for measured water chemistry parameters indicates distinct clustering of samples from disinfected and non-disinfected systems.

### Summary of metagenomic data set

Metagenomic analyses was used to assess the association between presence/absence of disinfectant residual with the structure and functional potential of the drinking water microbiome. A total of 41 drinking water samples were collected from DWDSs with (i.e., chlorine) from the United Kingdom (n=23), while those collected from the Netherlands (n=18) did not have a disinfectant residual. Quality trimming of raw metagenomic data resulted in the retention of 638 million paired-end reads. Co-assembly for each drinking water system was carried out by combining reads from individual sampling location within each drinking water system (Table 1). *De novo* co-assembly generated 0.04-1.81 million true scaffolds for each sampling location after discarding scaffolds shorter than 500bp and contaminant scaffolds (Table 1) with an N50 value ranged from 775 bp to 3300 bp. The proportion of quality trimmed reads mapping back to true scaffolds ranged from 67% to 99% (Table 1) across all samples.

**Table 1:**
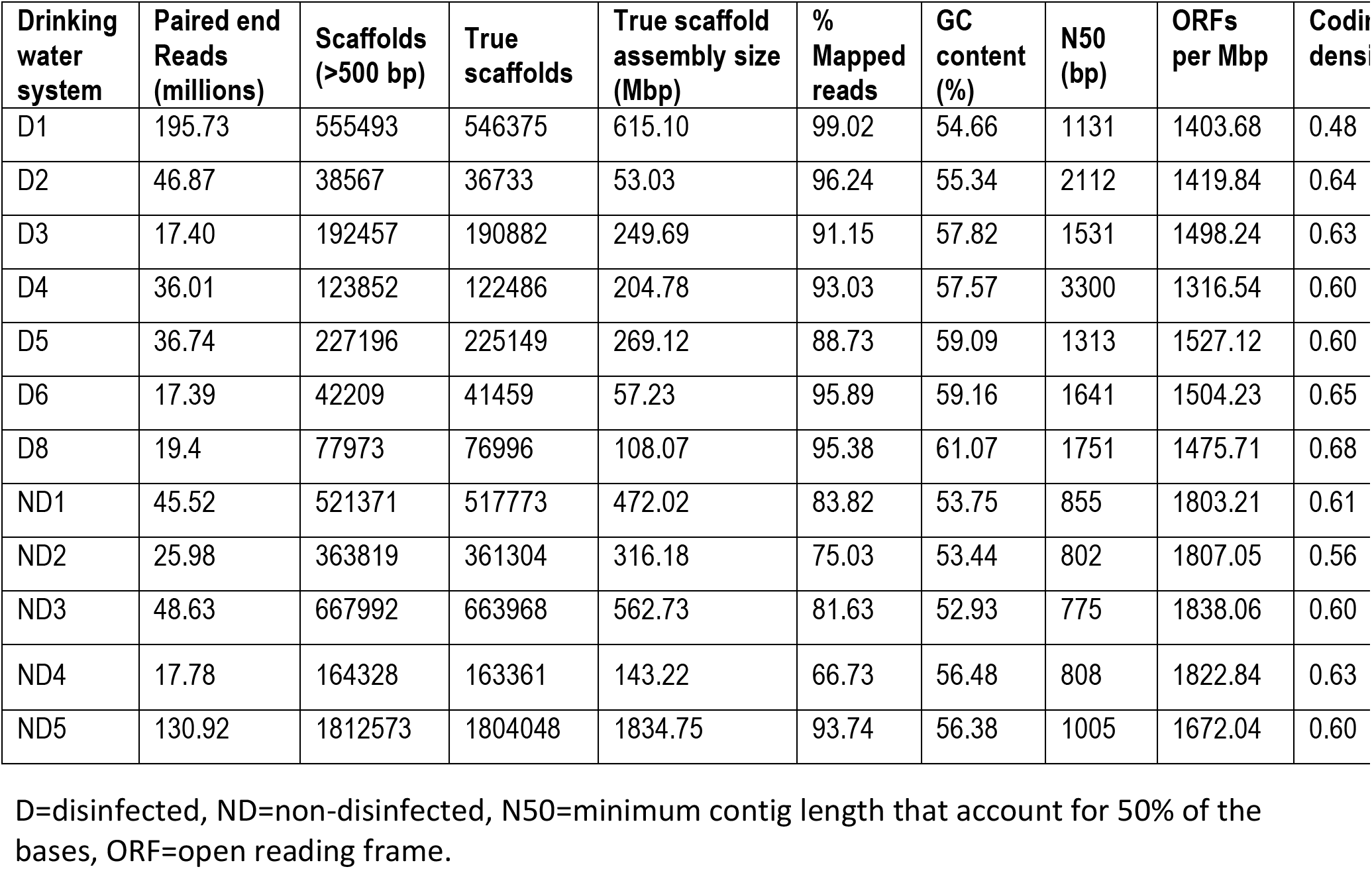
Sequencing and de novo co-assembly statistics for metagenomes from 12 drinking water systems.

### Non-disinfected systems are more diverse than disinfected systems

Non-disinfected systems were significantly (*p* < 0.0001) more diverse compared to systems that maintained a disinfectant residual (Figure *2A*) based on Nonpareil estimated diversity index^42^. This observation is consistent with previous comparisons of bulk water^69^ and biofilm^39^ samples from disinfected and non-disinfected systems. Lower diversity in disinfected systems is likely due to stronger selective pressure of the disinfectant residual as compared to that nutrient limitation in non-disinfected systems. As a result of the higher diversity in non-disinfected systems, the metagenomic sequencing for these samples provided significantly lower coverage of the sampled microbial community (Figure 2*B*) as compared to systems with a disinfectant residual (*p* < 0.0001).

**Figure 2:**
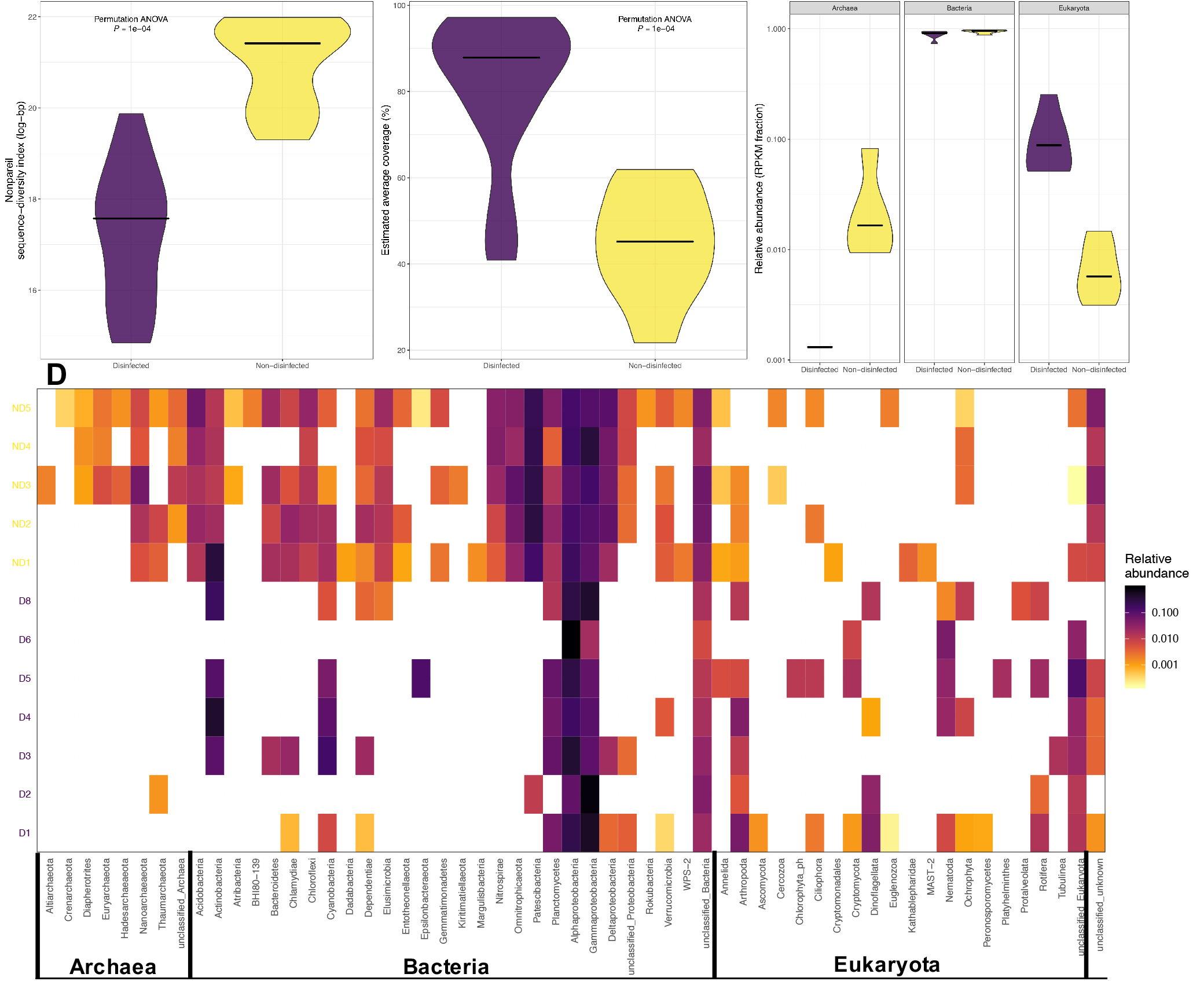
Comparison of (A) diversity and (B) coverage between disinfected and non-disinfected drinking water systems estimated using Nonpareil. (C) Comparison of relative abundance of bacterial, archaeal, and eukaryotic communities in drinking water systems with and without disinfectant residuals. (D) Log10 transformed relative abundance of different phyla (classes for phylum Proteobacteria) across sampling location for the bacteria, archaea, and eukaryota.

### Microbial community membership and structure is different between disinfected and non-disinfected systems

We used 2,872 small-subunit (SSU) rRNA genes (2742 genes > 100 bp) identified on the assembled scaffolds to determine community membership and structure across sampling locations (Supplementary file 1, Supplementary table 2). While bacteria were dominant members of the drinking water microbiome in both types of systems (2C, 2D), the relative abundance of archaea and eukaryota were dependent on the presence/absence of disinfectant residual (Figure 2C, 2E). Specifically, the relative abundance of eukaryota was higher in disinfected systems as compared to non-disinfected system (2C), while archaea were ubiquitous across non-disinfected samples (Figure 2C, E) they were only detected in a single disinfected sample (D2). Non-disinfected systems were taxonomically more diverse, with respect to bacteria and archaea, as compared to disinfected systems. Specifically, a total of 14 bacterial and 6 archaeal phyla were detected in one or more non-disinfected systems that were not detected in any of the disinfected systems. Several of these unique phyla, while not dominant in non-disinfected systems, were detected at relative abundances between 1-5% (e.g., *Nitrospirae, Nanoarchaeota*).

The bacterial community was dominated by *Proteobacteria*, in particular *Alphaproteobacteria* and *Gammaproteobacteria*, in both disinfected and non-disinfected systems with *Deltaproteobacteria* being much more prevalent and abundant in non-disinfected systems (Figure 2D). *Actinobacteria* were more abundant than *Proteobacteria* in two drinking water systems and constituted 44% and 33% of the community in systems D4 and ND1, respectively. Overall, the relative abundance of *Proteobacteria* was higher in disinfected systems, ranging from 28% to 90%, as compared to non-disinfected systems, ranging from 30% to 57%. *Patescibacteria* was the second most abundant phylum across non-disinfected systems, constituting 15% to 29% of the SSU rRNA genes, while they were only detected in one disinfected sample (D2) with a relative abundance of 1%. Within *Patescibacteria, Parcubacteria* were the most commonly detected phyla followed by *Microgenomatia* and *Gracilibacteria*.

The observed differences between disinfected and non-disinfected DWDS for bacteria and archaea are largely consistent with a previous meta-analyses of amplicon sequencing data from the 16S rRNA gene^69^. In contrast to bacteria and archaea, results from eukaryotes, which have not been systematically investigated in the drinking water microbiome, were surprising in terms of their higher relative abundance eukaryotic in disinfected as compared to non-disinfected systems. For instance, SSU rRNA genes associated *Nematoda* were detected in nearly every disinfected system, but were not detected in non-disinfected systems. Specifically, SSU rRNA genes from two free-living nematode genera, i.e. *Araeolaimida* and *Monhysterida*, were detected in five of the eight disinfected systems. Similarly, SSU rRNA genes from the phylum *Rotifera* were only detected in disinfected systems and were largely associated with the monogonont rotifers within the genera *Ploimida*. While the relative abundance of scaffolds determined to be of eukaryotic origin was higher in disinfected compared to non-disinfected systems, this does not mean that eukaryotes were proportionally larger part of the drinking water microbiome in disinfected compared to the non-disinfected systems. Genome sizes of picoeukaryotic microbes can be orders of magnitude larger than that of bacteria and archaea and vary significantly between picoeukaryotes themselves. Further, the higher overall diversity and lower sequencing coverage (Figure 1) could also have resulted in under sampling of the eukaryotic community in non-disinfected systems.

### Drinking water systems cluster at the nucleotide level based on presence/absence of disinfectant residuals

Samples (for read based analyses) and drinking water systems (for scaffold based analyses) clustered with each other based on the presence/absence of disinfectant residuals (Figure 3A and 3B) based on Mash distance estimates. We further evaluated the significance and explanatory power of measured water chemistry parameters in explaining the observed clustering between disinfected and non-disinfected systems. To do this, we initially performed BioEnv analyses to identify water chemistry parameters and their combinations that were highly correlated with observed Mash distances between samples (Supplementary Table 3). This identified chlorine as being strongly correlated with the Mash distances between samples (R=0.54, p<0.001) while the maximum correlation between water chemistry and Mash distances was observed for a combination of chlorine, phosphate, and TOC (R=0.62, p<0.001). We subsequently utilized dbRDA to independently determine the environmental/water chemistry variables most significantly associated with Mash distances between samples. While chlorine was identified as a significant variable (p<0.01), dbRDA identified conductivity (p<0.001) and DO (p<0.01) as significant variables. Finally, variance partitioning analyses was used to determine the proportion of variance in the Mash distance matrices explained by individual and combination of variables identified as significant by dbRDA (Table S5). This resulted in chlorine, conductivity, and DO individually explaining ~17%, 12%, and 1% of the variance in the Mash distance matrix, with ~60% of the variance unexplained by these three variables.

**Figure 3:**
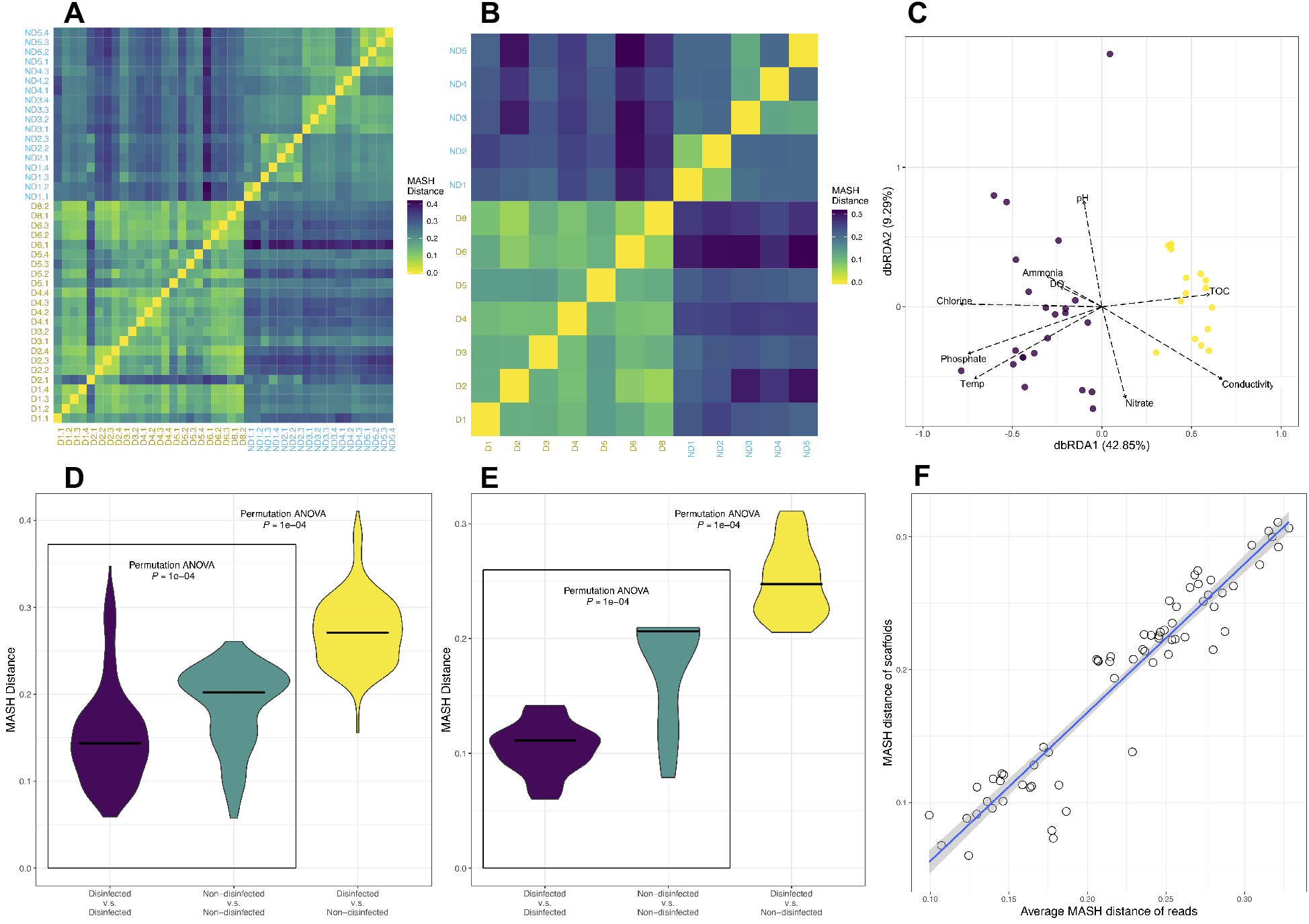
Comparison of nucleotide composition using paired reads each from each sample and true scaffolds in each drinking water system according to Mash distance. (A, B) Heatmaps based on pairwise Mash distances of reads and scaffolds. Heatmaps are colored according to Mash distance; yellow denotes a distance of 0. Labels on x- and y-axis are colored according to disinfection strategies. (C). NMDS clustering of read based Mash distances between samples with vectors representing water chemistry/environmental parameters implemented using dbRDA. (D, E) Violin plots indicating the distribution of pairwise Mash distances of reads and scaffolds. Plots are colored according to the system type for which pairwise comparisons were conducted. Purple denotes comparisons between disinfected samples, yellow denotes comparisons between non-disinfected samples, and green denotes comparisons between disinfected and non-disinfected samples. (F) Correlation between average Mash distances of reads across samples and Mash distances of scaffolds across sampling locations.

We further compared the distribution of Mash distances between drinking water metagenomes within disinfected, within non-disinfected, and between disinfected and non-disinfected systems. Mash distances between drinking water metagenomes from disinfected systems were significantly different (*p* < 0.0001) and exhibited a lower mean for disinfected as compared to non-disinfected systems. Further, the pairwise Mash-distances between disinfected and non-disinfected systems were significantly different and higher from those estimated within each category (i.e., disinfected or non-disinfected). This was consistent for both read- and scaffold-based analyses (Figure 3D, 3E). Finally, the average pairwise Mash distances estimated using reads (i.e., between samples) and scaffolds (i.e., between DWDSs) were highly correlated (Pearson’s *R* = 0.95, *P* < 0.05) (Figure 3C), indicating the *de novo* assembly process did not result in loss of information on factors driving the differences between disinfected and non-disinfected systems.

These analyses provide a few key insights. First, Mash distance-based (both read and scaffold based) clustering of samples occurs depending on presence and absence of disinfectant residual suggests that the microbial communities are more similar within each group (i.e., disinfected and non-disinfected) and dissimilar between the two groups (i.e., disinfected vs non-disinfected). Second, while disinfected and non-disinfected samples cluster distinctly from each other, disinfected systems exhibit lower nucleotide-level heterogeneity as compared to their non-disinfected systems indicating that the factors governing microbial community in disinfected systems likely impose stronger selective pressures on the microbial community as compared to those in non-disinfected systems. Third, non-disinfected systems exhibit greater diversity not only within a system (Figure 2) but also across systems as compared to disinfected systems. Despite the strong correlation between pairwise Mash distances of reads and scaffolds (Figure 3F), the median Mash distances for pairwise comparison of samples within each type of system (i.e., disinfected and non-disinfected) is higher for the scaffold-based analyses as compared to the read-based analyses. This is likely from the omission of low abundance microorganisms during *de novo* assembly and thus suggests that composition of medium-to-high abundance organisms are likely to be more variable between non-disinfected systems as compared to disinfected systems.

Finally, while the water chemistry and environmental parameters between disinfected and non-disinfected systems were distinct (Figure 1B), the parameters that most strongly correlated with Mash distances between samples were limited to a combination of chlorine, phosphate, and TOC for BioEnv analyses and chlorine, conductivity, and DO based on dbRDA. Both independent exploratory analyses consistently identified chlorine presence/absence and concentration as one of the key drivers of difference in microbial communities across the samples. Further, variance partition analyses indicated that ~17% of the variance in the Mash distance matrix was driven exclusively by chlorine; this make chlorine the most important parameter measured as part of this study in terms of differentiating between drinking water metagenomes. The significance of phosphate determined by BioEnv analyses is likely because chlorine and phosphate concentrations are inherently associated due to common use of the latter for corrosion control in DWDSs that maintain a chlorine residual^68^. Further, while it is unlikely that DO (identified as significant by dbRDA) directly affects microbial community composition (all DO concentrations were near or greater than saturation), it is possible that this may reflect the use of advanced oxidation process (e.g., ozonation) during drinking water treatment. Similarly, conductivity (identified as significant by dbRDA) is unlikely to directly influence the microbial community, but rather this may reflect the source water type and treatment processes being used for drinking water production. Specifically, source water derived from ground water sources or from reservoirs under the influence of ground water typically have much higher conductivities than those that rely on surface water supply. Similarly, chemicals used for softening and coagulation/flocculation processes may influence water conductivity. Thus, we speculate that the influence of conductivity may serve as surrogate for a combination of source water and treatment process. These analyses clearly identify chlorine as one of the major measured parameters driving the Mash distances between samples, followed by conductivity (potential surrogate for source water and treatment process). Further, the fact the major proportion of the variance remains unexplained suggests that additional aspects such as treatment process configuration, DWDS characteristics, and other water chemistry parameters which were not characterized/measured as part of this study also likely play a strong role in differentiating between microbial communities in disinfected and non-disinfected drinking water systems.

### Protein coding sequences cluster based on presence/absence of disinfectant residuals

A total of 8 million protein coding sequences were predicted and translated from true scaffolds, of which approximately 17 to 27% were annotated against KEGG database (Table S6). Consistent with the nucleotide-level analyses, samples clustered based on the presence and absence of disinfectant residual (Figure 4A, 4B, 4C) rather than by DWDS. Further, BioEnv analyses identified the combination of chlorine, phosphate, and ammonia as being strongly and significantly correlated (R =0.392, P<0.001) with Bray-Curtis distances between samples estimated using abundance (i.e., RPKM) of KOs (Table S7). Similar to nucleotide based analyses, chlorine presence/absence and concentration was the measured parameter more strongly and significantly associated with differences in functional potential between samples at the single parameter level (R=0.382, p<0.001). In contrast to nucleotide based analyses, conductivity and chlorine were the only two variables identified as significantly associated with Bray-Curtis distances between samples estimated using relative abundance of KO’s in samples using dbRDA (Table S8). Variance partitioning indicated that both conductivity and chlorine individually explained approximately 6.5% of the variance in Bray-Curtis distance matrix estimated using KO abundance. A comparison of the pairwise Mash distances within each group (i.e., disinfected, non-disinfected) and between them indicated that the diversity in functional potential was significantly different for both predicted protein coding-sequences and KEGG annotated proteins (*p* < 0.0001). The median value of Mash distances between the non-disinfected samples was greater than that for disinfected samples (Figure 4D, 4E) and the differences in Mash distances between two groups was larger than the distances within each group. And finally, despite the fact that only 17-27% of predicted proteins were annotated against the KEGG database, the Mash distances between metagenomes estimated using all predicted protein coding sequences and those that were annotated against the KEGG database were highly correlated (Pearson’s *R* ≈ 1.00, *P* < 0.05) (Figure 4F), suggesting that focusing on annotated proteins does not result in significant loss of information while performing direct comparisons between samples from disinfected and non-disinfected systems.

**Figure 4:**
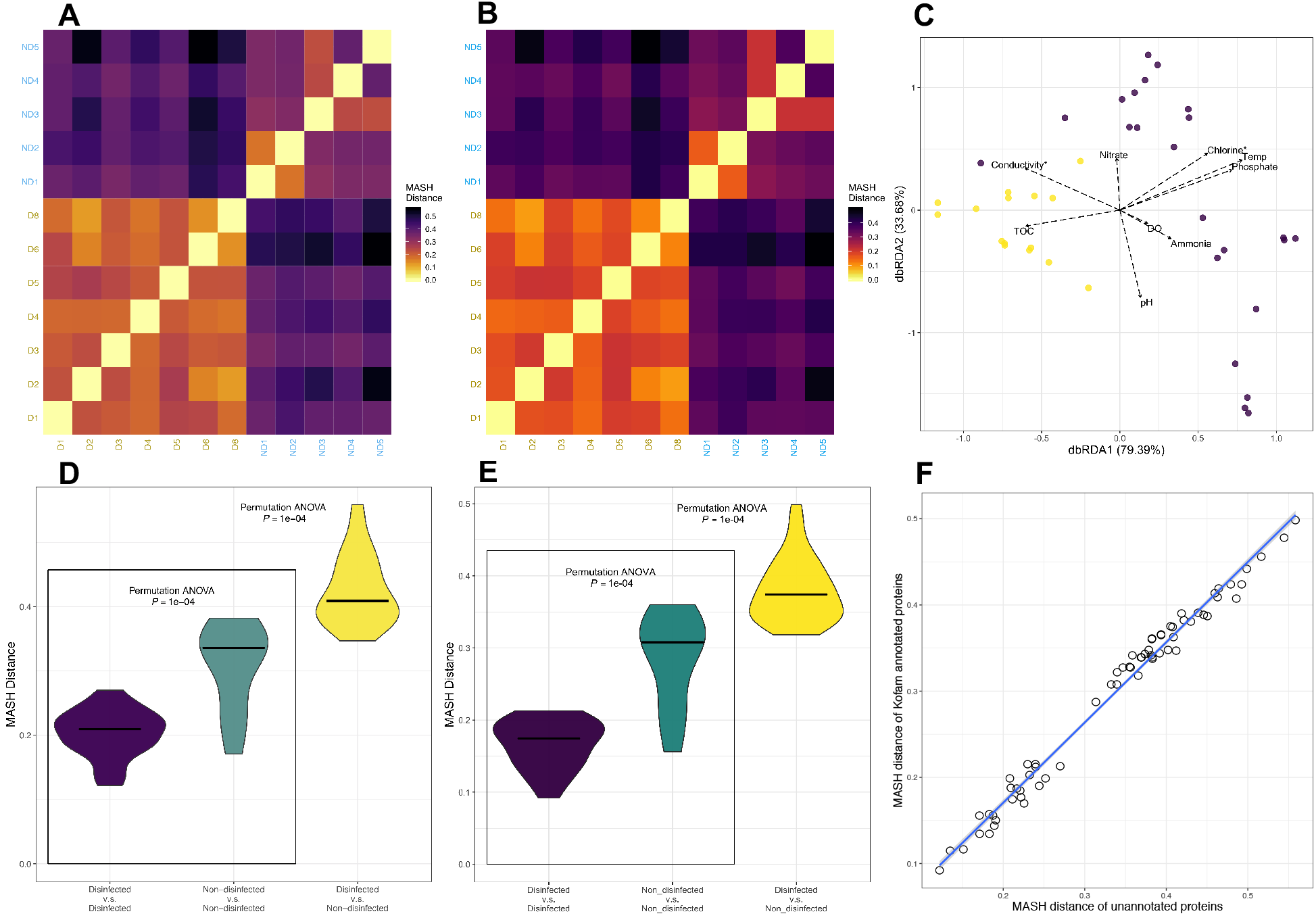
Comparison of functional potential among all and KEGG protein-coding amino acid sequences across sampling locations. This analysis estimates dissimilarity in amino acid composition of samples, similar to the nucleotide composition analyses presented earlier. (A, B) Heatmaps based on pairwise Mash distances of all protein coding sequences and Bray-Curtis distances using KO. Heatmaps are colored according to Mash/Bray-Curtis distance; yellow denotes a distance of 0. Labels on x- and y-axis are colored according to disinfection strategies; dark golden denotes samples with chlorine, while blue denotes samples without disinfectant residuals. C). NMDS clustering of using Bray-Curtis distances using KO abundances between samples with vectors representing water chemistry/environmental parameters implemented using dbRDA. Violin plots indicating the distribution of pairwise (D) Mash distances of all and (E) Bray-Curtis distances KEGG annotated proteins. Crossbars indicate the median value of Mash distances. (F) Correlation between pairwise Mash distances estimated using all and Bray-Curtis distances for KEGG annotated protein coding sequences.

These analyses based on protein coding sequencing provide several key insights. First, clustering of samples into disinfected and non-disinfected groups is consistent for both community composition (i.e., read-based nucleotide composition analyses) and functional potential, irrespective of the use of all predicted ORF’s and KEGG annotated protein sequences. Non-disinfected systems are significantly more heterogeneous across systems as compared to their disinfected counterparts. This suggests that selection pressures exerted within disinfected systems are not only evident at community structure/membership (Figure 3), but also evident at the community functional potential level. Further, consistent with microbial community composition, chlorine was also identified as one of the key measured parameters driving differences between samples based on functional potential using both BioEnv and dbRDA analyses. In contrast to TOC which was included in the BioEnv parameter combination for microbial community composition level analyses, ammonia was identified as part of the combination at the functional potential level. While the exact reason behind this difference cannot be ascertained in this study, this may likely be associated with the fact that non-disinfected systems are severely nitrogen limited as compared to disinfected systems, while both systems were likely not carbon limited. Similar to the nucleotide level analyses, both conductivity and chlorine were identified as significantly (p<0.01) associated with differences between samples, with variance partitioning analyses allocating equal amount of variation to both parameters (Table S9). As speculated above, if conductivity is considered a signal for source water and treatment process type, then the impact of these two parameters on the functional potential of microbial community is relatively similar to that of presence/absence of the disinfectant residual. Finally, the residuals from the variance partitioning analyses were noticeably larger (84%) for functional potential analyses as compared to the microbial community composition (60%), suggesting that the impact of unmeasured/uncharacterized factors/parameters on microbial community functional potential was significantly larger than their impact on community composition. While it cannot be ruled out, it is unlikely that the higher fraction of unexplained variation was due to only a proportion of ORFs being annotated; this is because Mash distances estimated using only KEGG annotated ORFs were highly correlated with those estimated using all predicted ORFs using suggesting little to minimal loss of discriminatory power while using only annotated proteins.

### Differentially abundant metabolic modules are consistent with microbial growth control strategies

A total of 7,281 KOs were identified in all samples with 5,922 remaining post-filtering based on scaffold coverage (>1x) and frequency of KO detection in each drinking water system (detected more than once) (Table S10). The 5,922 KO’s were further categorized into 540 KEGG modules and upon further filtering to remove KEGG modules with no more than one missing block and greater than equal to 50% completion, a total of 208 KEGG modules were retained (Table S11). Of these, a total of 57 KEGG modules exhibited significantly differential abundance between disinfected and non-disinfected samples (*p*-value < 0.005) (Table S12, S13). Modules associated with ribosomal synthesis, ribonucleotide biosynthesis, and RNA polymerase were ignored from further consideration. Similarly, modules most likely associated with plant metabolism (e.g., Crassulacean acid metabolism) were also ignored. This resulted in 29 and 22 KEGG modules that were more abundant in non-disinfected system and disinfected systems, respectively. These included modules associated with energy metabolism (disinfected, i.e. D=2, non-disinfected, i.e., ND=5), carbohydrate and lipid metabolism (D=11, ND=10), nucleotide and amino acid metabolism (D=5, ND=13), and secondary metabolism (D=4, ND=1).

Metabolic modules associated with polyamine biosynthesis, aromatics degradation, terpenoid biosynthesis, and fatty acid metabolism were significantly enriched in disinfected systems. Specifically, metabolic pathways associated with benzene (M00548) and benzoate (M00551) degradation to catechol and methyl catechol were highly enriched in disinfected systems. Further, eukaryota-associated metabolic modules such as terpenoid backbone biosynthesis (M00367) and modules associated with peroxisomal beta-oxidation of very long chain fatty acids (M00861) are likely to be enriched in the disinfected systems due to the higher relative abundance of eukaryota in samples collected from disinfected as compared to non-disinfected systems respectively. Further, modules related to *γ*-aminobutyrate (GABA) metabolism (M00136, M00027) were enriched in disinfected systems. The GABA shunt pathway converts glutamate to GABA using glutamate decarboxylase (GAD), followed by reversible conversion from *α*-ketoglutarate to succinate semialdehyde (SSA) through the activity of GABA transaminase (GABA-AT), and finally succinate is formed by succinate semialdehyde dehydrogenase (SSDH) activity. In contrast, the key metabolic modules enriched in non-disinfected systems were associated with carbon fixation and methane metabolism (M00377, M00620, and M00422) and nitrogen fixation (M00175) (Table S13). The differentially abundant carbon fixation modules included the Wood-Ljungdahl pathway, Acetyl-CoA pathway, and the incomplete reductive citrate cycle. These pathways can fix carbon dioxide to produce acetyl-CoA which can then be converted to other necessary biosynthetic intermediates of the carbon metabolism^70, 71^.

The enrichment of carbon and nitrogen fixation modules in non-disinfected systems is consistent with nutrient limitation as the strategy for microbial growth control in non-disinfected drinking water systems. While the measured total organic carbon concentrations in non-disinfected systems did not indicate carbon limited conditions, DWTP’s supplying water to non-disinfected DWDSs typically achieve far superior levels of removal of assimilable organic carbon (AOC)^28^. Similarly, the nitrogen availability in the form of ammonia was consistently zero for non-disinfected systems compared to disinfected systems which has residual ammonia concentrations ranged from 0.01-0.15 mg/l of ammonia-nitrogen. In contrast, the enrichment of KEGG modules associated with GABA metabolism in disinfected systems suggests the potential importance of stress protection and utilization of microbial decay products. Previous studies have shown that GABA metabolism is associated with bacterial survival under various types of environmental stresses, including oxidative stress, acidic stress, and osmotic stress^72–75^. Meanwhile, GABA can also play a significant role in nitrogen metabolism of bacteria. For instance, putrescine formed due to the breakdown of amino acids potentially from decaying biomass, can be converted to GABA (M00136) and finally metabolized via GABA shunt pathway^74^. The enrichment of GABA metabolism in disinfected systems may thus be associated with greater protection against disinfectant stress and by allowing access to decay products from inactivated cells.

### Average genome size differences between disinfected and non-disinfected system vary between read-based and MAG-based analyses

We further investigated differences in genome sizes between disinfected and non-disinfected systems. Genome sizes can be indicative of the metabolic capacity of microorganisms^76^ and thus provide insights in the whether the presence/absence of disinfectants selects for organisms with larger or smaller metabolic repertoire^77^ in comparison to organisms detected in non-disinfected systems. Average genomes size estimates from disinfected systems were significantly larger than those from non-disinfected systems based on MicrobeCensus estimates using entire metagenomic data (Figure 5A); this was consistent even when reads mapping to phyla known to have smaller genomes (e.g., *Patescibacteria*) were selectively removed from the data set (Figure 5B). This suggests that microorganisms in disinfected systems may be metabolically more diverse than their counterparts from non-disinfected systems. Nonetheless, these results were not consistent when compared with estimated genome sizes of MAGs recovered as part of this study. Specifically, we recovered a total of 115 dereplicated MAGs with completeness >50% and redundancy <10% (Table S14). These 115 MAGS were binned into four categories based on the detection or non-detection in disinfected samples. Specifically, MAGs were binned in the four groups (i.e., both, D-only, ND-only, and other) based on genome coverage and detection frequency criteria outlined in the materials and methods section (see MAG-level analyses) (Table S15). This resulted in 9, 16, 41, and 49 MAGs were categorized as both, D-only, ND-only, and other (Figure 5C, 5D) (Table S14). In contrast to read-based estimates of average genome size, MAG-based genome size estimates were not significantly different between the three key categories (Both=4.4±0.77Mbp, D-only=3.22±0.81Mbp, ND-only=3.48±1.22Mbp) (Figure 5E). Yet, the ND-only category consisted of several smaller genomes (n=17) compared to the D category. The lack of genome size differences between disinfected and non-disinfected samples based on MAG-based analyses compared to metagenome-level read-based analyses may be due to the proportion of read-based data represented by the MAGs. Specifically, while 60-90%of the reads from disinfected systems mapped to the 115 MAGs with the mapping rate from non-disinfected systems averaging around 50% (Figure 5F). Thus, it is likely that the metagenomic assembly and binning process may have resulted in suboptimal recovery of smaller genomes from non-disinfected sample which eliminates the signal in genome size differences observed at the metagenome level.

**Figure 5:**
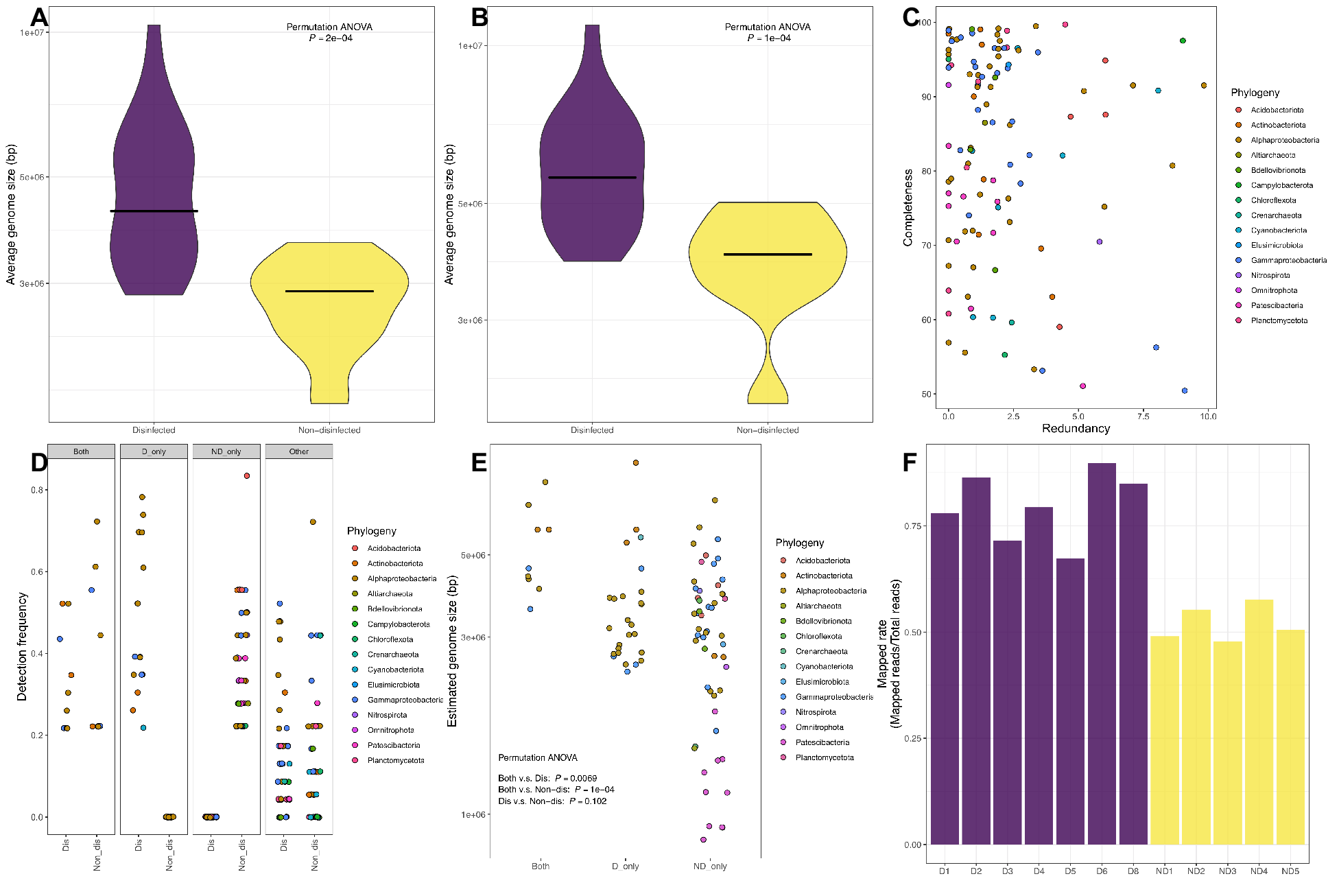
Violin plots indicating the genome size estimated by MicrobeCensus (a) before and (b) after Patescibacteria removal suggest average genome sizes in disinfected systems are larger than those in non-disinfected systems. (C) The 115 MAGs assembled with >50% completeness and <10% redundancy were categorized into (D) four groups based on their detection frequency in disinfected and non-disinfected systems. (E) While the estimated genome sizes of MAGs in D_only, ND_only, and Both categories were not significantly different, the ND_only category consisted of large number of smaller genomes. (F) Barplot indicating the proportion of reads mapped to 115 genomes across samples. Purple and yellow denotes samples from systems with and without a disinfectant residual, respectively.

### Metabolic capacities differ between metagenome assembled genomes from disinfected and non-disinfected systems

Clustering of MAGs (Figure 6A) based on presence/absence of KEGG metabolic modules was largely driven by phylogenetic placement of MAGs, rather than their classifications into groups based on the detection frequencies in disinfected and non-disinfected systems (Figure 6B). Further, there was insufficient representation of MAGs from D-only/ND-only categories across all phylogenetic clusters (e.g., at the species or genus level) to allow for direct comparisons of metabolic potential of closely related MAGs exclusively frequent in disinfected and non-disinfected systems. Nonetheless, there were seven and five high quality (completeness > 90%, redundancy <10%) alphaproteobacterial MAGs that were exclusively frequent in disinfected (average detection frequency in disinfected =55%) and non-disinfected systems (average detection frequency in non-disinfected=29%) (Figure 6A). Thus, we focused metabolic module comparisons between these 12 MAGs only. We evaluated differences in metabolic capacity of these MAGs by (1) considering all KEGG modules ≥75% complete within MAGs to be present in them and (2) all modules present in more than half of the high-quality MAGs within each category to be present within each category (Figure 6C, Table S16). We subsequently confirmed the presence/absence of genes within key metabolic modules using KO-level annotation for these 12 MAGs (Table S17).

**Figure 6:**
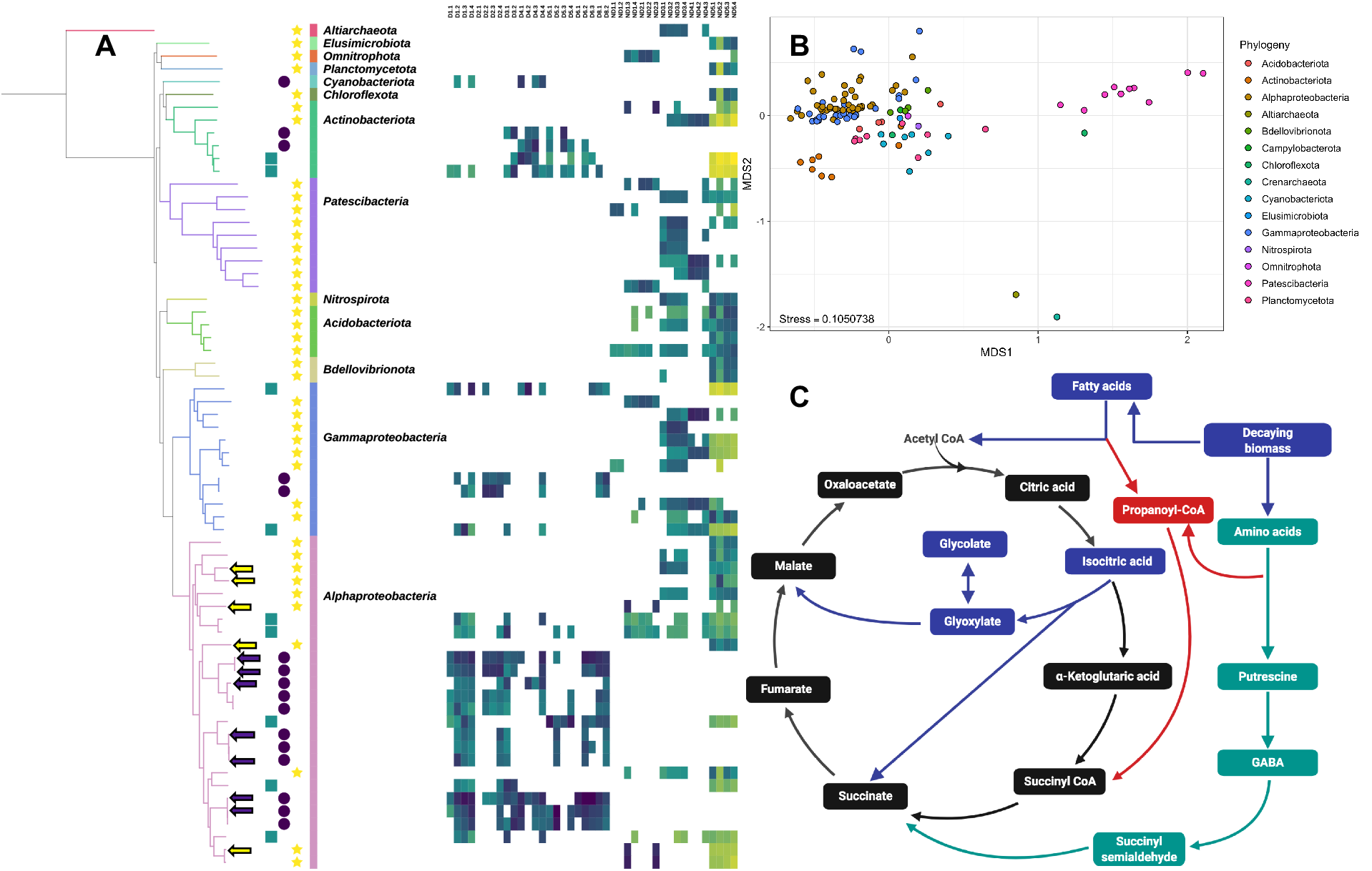
(A) Phylogenomic tree of 66 MAGs classified as D-only (purple circles), ND-only (yellow stars), and both (teal squares) constructed using 48 ribosomal proteins along and their relative abundance (RPKM) in the samples collected from disinfected and non-disinfected systems. RPKM’s for MAGs are only reported for samples where 25% of the nucleotides in a MAG were covered by at least one read. (B) Clustering of all MAGs based on their clustering metabolic potential (i.e., completeness of KEGG modules) was primarily drive by phylogeny. (C) The metabolic modules identified as differentially abundant in disinfected systems using metagenome level analyses (Table S12) are shown using teal arrows and squares and those more prevalent in high quality alphaproteobacterial MAGs from D-only (purple arrows - Figure 6A) compared to those from ND-only category (yellow arrows - Figure 6A) are shown using blue arrows and boxes, while red arrows and boxes indicates modules identified as more prevalent in D-only systems using both metagenome and MAG level analyses.

The metabolic module associated with the glyoxylate cycle (M00012) was present in 86% of the MAGs in the D-only category while being only partially complete in most of the ND-only MAGs. Specifically, isocitrate lyase (*aceA:* K01637) and malate synthase (*aceB*: K01638), two key genes involved in the glyoxylate cycle, were present in 40% and 100% of the MAGs from D-only, respectively and both genes were absent in all ND-only MAGs included in this analysis. The glyoxylate shunt is associated with use of non-carbohydrate carbon sources (i.e., via gluconeogensis), such as break down products from lipids, fatty acids etc^78^. The likely benefit of the glyoxylate shunt and associated use of lipids and fatty acids as carbon source is further supported by the fact that KEGG module associated with propanoyl-coA metabolism (M00741) was complete in 6/7 as compared to 2/5 MAGs from the D-only and ND-only categories. This metabolic module is associated with the conversion of propanoyl-coA, a toxic byproduct of fatty and amino acid degradation, to succinyl-coA. High biomass turnover rates, due to disinfectant induced microbial inactivation, may result in resource pools enriched in microbial decay products thus allowing a significant advantage for microorganisms capable of necrotrophic growth^79^ aided by the glyoxylate cycle. Thus, it is feasible that the ability to utilize microbial decay products may provide a distinct advantage to microorganisms inhabiting disinfected DWDSs.

The glyoxylate shunt may provide additional benefits for microorganisms subject to disinfectant stress via enhanced fitness to oxidative stress^78^ and enhanced persistence when challenged with other chemical stressors (e.g., antibiotics)^80^. In contrast to module level analyses at the metagenome level where carbon fixation capacity was significantly more abundant in non-disinfected as compared to disinfected systems, the alphaproteobacterial MAGs from D-only systems harbored the capacity for carbon fixation via the Calvin-Benson-Bassham cycle (M00165, M00166, M00167) while this capacity was mostly absent from MAGs in the ND-only category. Nonetheless, these MAG-based analyses are limited in phylogenetic scope and does not weigh the importance of MAGs to their respective systems based on their relative abundance. Hence, we suggest that metagenome-level analyses should take precedence over findings at the MAG level when they conflict. While the glyoxylate shunt was not identified as a significantly enriched in the disinfected systems at the metagenome level analyses, the GABA shunt (metagenome level analyses) and glyoxylate shunt (MAG level analyses) may both be involved in use of non-carbohydrate carbon sources suggesting that re-use of microbial decay products may indeed be a key bacterial trait that allows for persistence in disinfected drinking water systems. Further lending support to this is that that propanoyl-coA metabolism was identified as significantly enriched in disinfected systems compared to non-disinfected systems using both metagenome-level and MAG-level analyses. Interestingly, only one metabolic module was identified as being more than twice as prevalent in alphaproteobacterial MAGs from ND-only systems compared to those from D-only systems (i.e., M00156: cbb3-type Cytochrome C oxidase). The greater metabolic capacity of alphaproteobacterial D-only MAGs compared to ND-only MAGs was also confirmed at the KO-level by evaluating the presence/absence of KO’s in the D-only and ND-only category MAGs. Specifically, while only 8 KOs were twice or more as prevalent in ND-only MAGs compared to D-only MAGs, the total KOs that were twice or more as prevalent in D-only MAGs was 109. This supports the conclusion that metabolic repertoire of alphaproteobacterial D-only MAGs is significantly larger than that of ND-only MAGs. Notable among the genes that were twice as frequent in D-only MAGs compared ND-only MAGs included those involve in SOS-response mediated mutagenesis involving trans-lesion synthesis (i.e., *imuA*: K14160, *imuB*: K14161, and *dnaE2*: K14162)^81^, glyoxylate reductase (*gyaR*: K00015) which may be likely involved in regulating glyoxylate concentrations, and vitamin B12 transporter (*btuB*: K16092). SOS response is typically activated in response to significant cellular accumulation of damaged DNA^82^ and *imuA*and *imuB* co-expression with *dnaE2* has been shown to be responsive to UV damage ^81^. Thus, the higher prevalence of SOS response related genes in D_only MAGs may be associated with the DNA damage caused by disinfectants. Further, the ability to synthesize vitamin B12, an essential co-factor, is limited to certain bacteria and archaea and thus the ability to uptake vitamin B12 from the environment is essential for growth^83^. The higher abundance of vitamin B12 transporters is consistent with metagenome level observations that the microbial community in disinfected systems rely more on scavenging from the environment as compared to non-disinfected systems.

## CONCLUSIONS

To our knowledge, this is the first study to provide metagenomic insights into differences in structure and functional potential of drinking water microbiomes across full-scale drinking water systems that rely on disinfection (i.e., disinfected) or nutrient limitation (i.e., non-disinfected) to manage microbial growth. Understanding the microbial implications of these two microbial growth control strategies is essential to not only develop a better understanding of ecological and metabolic traits guiding community level processes in these system, but is also critical for providing a community-level context to the microbiological safety in either type of drinking water system. In this study, we show that disinfection exhibits consistent, systematic, and significant association with drinking water microbiome at the membership, structure, and functional potential at the metagenome and MAG levels, irrespective of the drinking water system under consideration (e.g., source water type, treatment process, etc.). In doing so, we also identify key metabolic traits associated with carbon and nitrogen metabolism that are over represented in bacteria in disinfected systems compared to non-disinfected systems. This suggests that the influence and efficacy of disinfection on the drinking water microbiome may not simply be associated with differential disinfection resistance^84^, but may also expand to other metabolic traits that include the use of carbon and nitrogen sources made available via microbial inactivation and its regulation. It is important to note that while the impact of disinfection on microbial community structure and functional potential is clear, the metabolic traits identified in this study provide a hypothesis to support future experimental work that will be required to validate the findings of this study.

## ACKNOWLEDGEMENTS

ZD was supported by the Lord Kelvin Adam Smith Scholarship at the University of Glasgow. MS was supported by the College of Engineering at Northeastern University. This work was funded by Engineering and Physical Science Research Council (EP/M016811/1) and the National Science Foundation (NSF-CBET 1749530). UZI is supported by NERC Independent Research Fellowship (NERC NE/L011956/1). The authors are grateful to Prof. Karthik Anantharaman for providing helpful critiques of the manuscript.

