## Supplemental tables for "Disinfection exhibits systematic impacts on the drinking water microbiome"

**Table S1:** Summary of water quality parameters measured for the samples collected as part of this study. (*NM = not measured)

| **Sample** | **DWS** | **Type** | **Country** | **Temp** | **pH** | **Conductivity** | **DO** | **Total Chlorine** | **Phosphate** | **TOC** | **Ammonia** | **Nitrate** |
| --- | --- | --- | --- | --- | --- | --- | --- | --- | --- | --- | --- | --- |
|  |  |  |  | **(ºC)** |  | **(mS/cm)** | **(mg/l)** | **(mg Cl_2_/l)** | **(mg PO_4_^3-^/l)** | **(mg/l)** | **(mg N/l)** | **(mg N/l)** |
| D1.1 | D1 | Dis | UK | 16.60 | 8.54 | 128.90 | 4.99 | 0.36 | 1.09 | 2.29 | 0.02 | 0.60 |
| D1.2 | D1 | Dis | UK | 18.30 | 8.51 | 116.50 | 5.07 | 0.31 | 0.98 | 2.15 | 0.04 | 0.48 |
| D1.3 | D1 | Dis | UK | 16.60 | 8.47 | 113.00 | 5.41 | 0.10 | 1.12 | 1.94 | 0.03 | 0.42 |
| D1.4 | D1 | Dis | UK | 18.30 | 8.39 | 114.50 | 5.54 | 0.13 | 1.15 | 2.29 | 0.03 | 0.49 |
| D2.1 | D2 | Dis | UK | 13.30 | 8.39 | 65.50 | 10.76 | 0.73 | 2.44 | 2.64 | 0.08 | 0.21 |
| D2.2 | D2 | Dis | UK | 12.50 | 8.45 | 65.00 | 10.05 | 0.42 | 2.06 | 2.62 | 0.08 | 0.17 |
| D2.3 | D2 | Dis | UK | 12.30 | 8.42 | 66.00 | 10.10 | 0.43 | 2.10 | 2.72 | 0.08 | 0.37 |
| D2.4 | D2 | Dis | UK | 14.50 | 8.33 | 68.70 | 11.36 | 0.33 | 2.07 | 2.44 | 0.07 | 0.25 |
| D3.1 | D3 | Dis | UK | 19.90 | 7.88 | 144.75 | 10.37 | 0.16 | 1.23 | 1.39 | 0.00 | 1.49 |
| D3.2 | D3 | Dis | UK | 10.60 | 7.54 | 148.25 | 11.90 | 0.26 | 1.82 | 1.91 | 0.00 | 1.53 |
| D4.1 | D4 | Dis | UK | 15.10 | 7.56 | 313.00 | 8.92 | 0.66 | 1.97 | 1.80 | 0.00 | 1.10 |
| D4.2 | D4 | Dis | UK | 17.40 | 7.67 | 312.50 | 9.46 | 0.50 | 2.00 | 1.60 | 0.00 | 1.01 |
| D4.3 | D4 | Dis | UK | 18.30 | 7.95 | 311.50 | 9.17 | 0.60 | 2.12 | 1.63 | 0.00 | 1.07 |
| D4.4 | D4 | Dis | UK | 17.90 | 7.93 | 306.00 | 9.59 | 0.36 | 1.96 | 1.58 | 0.00 | 1.00 |
| D5.1 | D5 | Dis | UK | 19.20 | 7.49 | 666.00 | 8.11 | 0.63 | 4.94 | 2.32 | 0.14 | 6.07 |
| D5.2 | D5 | Dis | UK | 18.20 | 7.54 | 683.50 | 13.26 | 0.42 | 0.85 | 2.24 | 0.00 | 6.02 |
| D5.3 | D5 | Dis | UK | 21.10 | 7.92 | 652.00 | 8.86 | 0.36 | 4.67 | 3.61 | 0.13 | 5.68 |
| D5.4 | D5 | Dis | UK | 21.60 | 7.47 | 653.00 | 8.09 | 0.42 | 4.77 | 2.28 | 0.00 | 5.82 |
| D6.1 | D6 | Dis | UK | 17.90 | 7.88 | 216.40 | 9.68 | 0.38 | 3.06 | 1.00 | 0.00 | 1.70 |
| D6.2 | D6 | Dis | UK | 19.40 | 8.08 | 212.60 | 9.13 | 0.12 | 3.05 | 0.94 | 0.00 | 1.64 |
| D6.3 | D6 | Dis | UK | 19.70 | 8.02 | 227.65 | 9.33 | 0.26 | 3.09 | 1.08 | 0.00 | 1.73 |
| D8.1 | D8 | Dis | UK | 15.10 | 8.74 | 103.35 | 10.90 | 0.41 | 2.48 | 1.59 | 0.01 | 0.27 |
| D8.2 | D8 | Dis | UK | 12.65 | 8.50 | 101.15 | 10.93 | 0.11 | 2.37 | 1.48 | 0.01 | 0.20 |
| ND1.1 | ND1 | NonDis | NL | 11.40 | 8.25 | 542.00 | 8.98 | 0.00 | 0.03 | 1.01 | 0.00 | 0.62 |
| ND1.2 | ND1 | NonDis | NL | 11.80 | 8.23 | 546.00 | 8.74 | 0.00 | 0.03 | 1.01 | 0.00 | 0.60 |
| ND1.3 | ND1 | NonDis | NL | 12.00 | 8.24 | 546.00 | 8.70 | 0.00 | 0.03 | 1.09 | 0.00 | 0.61 |
| ND1.4 | ND1 | NonDis | NL | 14.80 | NM | NM | NM | 0.00 | 0.03 | 1.04 | 0.00 | 0.55 |
| ND2.1 | ND2 | NonDis | NL | 12.40 | 8.11 | 467.00 | 8.41 | 0.00 | 0.00 | 2.80 | 0.00 | 0.99 |
| ND2.2 | ND2 | NonDis | NL | 11.80 | 8.13 | 472.00 | 8.33 | 0.00 | 0.03 | 2.72 | 0.00 | 0.91 |
| ND2.3 | ND2 | NonDis | NL | NM | NM | NM | NM | 0.00 | 0.00 | 2.63 | 0.00 | 0.93 |
| ND3.1 | ND3 | NonDis | NL | 11.70 | 8.09 | 619.00 | 8.50 | 0.00 | 0.00 | 5.60 | 0.00 | 2.78 |
| ND3.2 | ND3 | NonDis | NL | 11.30 | 8.06 | 599.00 | 8.50 | 0.00 | 0.00 | 5.50 | 0.00 | 2.84 |
| ND3.3 | ND3 | NonDis | NL | 10.70 | 7.97 | 618.00 | 9.10 | 0.00 | 0.00 | 5.40 | 0.00 | 2.76 |
| ND3.4 | ND3 | NonDis | NL | 10.10 | 8.06 | 614.00 | 8.70 | 0.00 | 0.00 | 5.40 | 0.00 | 2.98 |
| ND4.1 | ND4 | NonDis | NL | 11.90 | 8.37 | 322.00 | 10.20 | 0.00 | 0.00 | 1.00 | 0.00 | 0.48 |
| ND4.2 | ND4 | NonDis | NL | 11.40 | 8.28 | 336.00 | 10.30 | 0.00 | 0.00 | 1.10 | 0.00 | 0.53 |
| ND4.3 | ND4 | NonDis | NL | 10.80 | 8.38 | 334.00 | 9.70 | 0.00 | 0.00 | 1.00 | 0.00 | 0.48 |
| ND5.1 | ND5 | NonDis | NL | 11.60 | 7.59 | 507.00 | 8.60 | 0.00 | 0.00 | 3.90 | 0.00 | 2.59 |
| ND5.2 | ND5 | NonDis | NL | 10.30 | 8.01 | 527.00 | 9.30 | 0.00 | 0.00 | 3.80 | 0.00 | 0.96 |
| ND5.3 | ND5 | NonDis | NL | 10.70 | 7.61 | 482.00 | 8.70 | 0.00 | 0.00 | 4.20 | 0.00 | 2.79 |
| ND5.4 | ND5 | NonDis | NL | 9.50 | 7.53 | 506.00 | 7.50 | 0.00 | 0.00 | 3.20 | 0.00 | 2.80 |

**Table S2:** SILVA database classification of SSU rRNA genes retrieved from assembled scaffolds. Scaffold names indicate DWS number, scaffold number, and the coordinates of the SSU rRNA gene on the scaffold. (See separate excel spreadsheet).

**Table S3:** BioEnv analyses in vegan package to determine the subset of variables significantly correlated with community similarities. This determines the Spearman’s correlation between Euclidean distances of scaled environmental variables with the Mash distances estimated using metagenomic reads.

| **Number of parameters** | **Parameter combination** | **Spearman's correlation** | **p value** |
| --- | --- | --- | --- |
| 1 | Chlorine | 0.54 | 0.001 |
| 2 | Chlorine + Phosphate | 0.58 | 0.001 |
| 3 | Chlorine + Phosphate + TOC | 0.62 | 0.001 |
| 4 | Temp + Chlorine + Phosphate + TOC | 0.61 | 0.001 |
| 5 | Temp + Conductivity + Chlorine + Phosphate + TOC | 0.61 | 0.001 |
| 6 | Temp + pH + Conductivity + Chlorine + Phosphate + TOC | 0.56 | 0.001 |
| 7 | Temp + pH + Conductivity + Chlorine + Phosphate + TOC + Ammonia | 0.51 | 0.001 |
| 8 | Temp + pH + Conductivity + Chlorine + Phosphate + TOC + Ammonia + Nitrate | 0.47 | 0.001 |
| 9 | Temp + pH + Conductivity + DO + Chlorine + Phosphate + TOC + Ammonia + Nitrate | 0.40 | 0.001 |

**Table S4:** Distance based Redundancy Analysis using Mash distance matrix generated using pairwise Mash distances between samples estimated using metagenomic reads.

Permutation test for dbrda under reduced model

Marginal effects of terms

Permutation: free

Number of permutations: 9999

**Model**: dbrda(formula = Mash_distance.dist ~ pH + Conductivity + Chlorine + Temp + DO + Phosphate + TOC + Ammonia + Nitrate, data = meta_table[, selected_environmental_data, drop = F])

| **Variables** | **Degrees of Freedom** | **Sum of Squares** | **F** | **Pr (>F)** | **Significance** |
| --- | --- | --- | --- | --- | --- |
| pH | 1 | 0.028051 | 2.5851 | 0.0518 | . |
| Conductivity | 1 | 0.061962 | 5.7103 | 0.0029 | ** |
| Chlorine | 1 | 0.036368 | 3.3516 | 0.021 | * |
| Temperature | 1 | 0.026548 | 2.4466 | 0.0593 | . |
| Dissolved oxygen | 1 | 0.033352 | 3.0737 | 0.0313 | * |
| Phosphate | 1 | 0.013299 | 1.2256 | 0.2787 |  |
| Temperature | 1 | 0.020983 | 1.9338 | 0.1078 |  |
| Ammonia | 1 | 0.013206 | 1.217 | 0.2704 |  |
| Nitrate | 1 | 0.023434 | 2.1596 | 0.0836 | . |
| Residual | 28 | 0.303824 |  |  |  |

Signif. codes: 0 ‘***’ 0.001 ‘**’ 0.01 ‘*’ 0.05 ‘.’ 0.1 ‘ ’ 1

**Table S5:** Variance Partition analyses using water chemistry/environmental parameters identified as significant being significantly associated with read-based Mash distances by dbRDA analyses.

Command used:

Mash_var<-varpart(Mash.dist, ~Chlorine, ~Conductivity, ~DO, data = metadata)

No. of explanatory tables: 3
Total variation (SS): 0.96648
No. of observations: 38

X1 = Chlorine, X2 = Conductivity, X3 = Dissolved oxygen (DO)

| **Variables and their combinations** | **Degrees of freedom** | **R square** | **Adjusted R square** | **Testable** |
| --- | --- | --- | --- | --- |
| [a+d+f+g] = X1 | 1 | 0.29 | 0.27 | TRUE |
| [b+d+e+g] = X2 | 1 | 0.23 | 0.21 | TRUE |
| [c+e+f+g] = X3 | 1 | 0.04 | 0.02 | TRUE |
| [a+b+d+e+f+g] = X1+X2 | 2 | 0.42 | 0.38 | TRUE |
| [a+c+d+e+f+g] = X1+X3 | 2 | 0.31 | 0.28 | TRUE |
| [b+c+d+e+f+g] = X2+X3 | 2 | 0.27 | 0.23 | TRUE |
| [a+b+c+d+e+f+g] = All | 3 | 0.45 | 0.40 | TRUE |
| Individual fractions |  |  |  |  |
| [a] = X1\|X2+X3 | 1 |  | 0.17 | TRUE |
| [b] = X2\|X1+X3 | 1 |  | 0.12 | TRUE |
| [c] = X3\|X1+X2 | 1 |  | 0.01 | TRUE |
| [d] | 0 |  | 0.09 | FALSE |
| [e] | 0 |  | -0.01 | FALSE |
| [f] | 0 |  | 0.01 | FALSE |
| [g] | 0 |  | 0.00 | FALSE |
| [h] = Residuals |  |  | 0.60 | FALSE |
| Controlling 1 table X |  |  |  |  |
| [a+d] = X1\|X3 | 1 |  | 0.26 | TRUE |
| [a+f] = X1\|X2 | 1 |  | 0.17 | TRUE |
| [b+d] = X2\|X3 | 1 |  | 0.21 | TRUE |
| [b+e] = X2\|X1 | 1 |  | 0.12 | TRUE |
| [c+e] = X3\|X1 | 1 |  | 0.01 | TRUE |
| [c+f] = X3\|X2 | 1 |  | 0.02 | TRUE |

**Table S6:** Number of predicted open reading frames (ORFs) for each metagenome co-assembly and number annotated against the KEGG database using Kofamscan.

| **DWS** | **Total predicted ORF's** | **ORF's annotated using KEGG database** | **Percent ORF's identified using KEGG database** |
| --- | --- | --- | --- |
| D1 | 863400 | 149048 | 17.26 |
| D2 | 75290 | 20440 | 27.15 |
| D3 | 374097 | 82883 | 22.16 |
| D4 | 269598 | 61517 | 22.82 |
| D5 | 410971 | 85531 | 20.81 |
| D6 | 86083 | 21343 | 24.79 |
| D8 | 159482 | 42639 | 26.74 |
| ND1 | 851157 | 172437 | 20.26 |
| ND2 | 571361 | 102030 | 17.86 |
| ND3 | 1034340 | 205383 | 19.86 |
| ND4 | 261064 | 56331 | 21.58 |
| ND5 | 3067778 | 606300 | 19.76 |

**Table S7:** BioEnv analyses in vegan package to determine the subset of variables significantly correlated with similarities in functional potential of community estimates using KEGG annotation. This determines the Spearman’s correlation between Euclidean distances of scaled environmental variables with Bray Curtis distance matrix generated from RPKM of KO detected in samples.

| **Number of parameters** | **Parameter combination** | **Spearman's correlation** | **p value** |
| --- | --- | --- | --- |
| 1 | Chlorine | 0.382 | 0.001 |
| 2 | Chlorine + Ammonia | 0.381 | 0.001 |
| 3 | Chlorine + Phosphate + Ammonia | 0.392 | 0.001 |
| 4 | Conductivity + Chlorine + Phosphate + Ammonia | 0.369 | 0.001 |
| 5 | Temp + Conductivity + Chlorine + Phosphate + Ammonia | 0.361 | 0.001 |
| 6 | Temp + Conductivity + Chlorine + Phosphate + Ammonia + Nitrate | 0.346 | 0.001 |
| 7 | Temp + pH + Conductivity + Chlorine + Phosphate + Ammonia + Nitrate | 0.320 | 0.001 |

**Table S8:** Distance based Redundancy Analysis using pairwise Bray-Curtis distances between samples estimated using from RPKM of KO detected in samples.

Permutation test for dbrda under reduced model

Marginal effects of terms

Permutation: free

Number of permutations: 9999

**Model:** dbrda(formula = BC_distance ~ pH + Conductivity + Chlorine + Temp + DO + Phosphate + TOC + Ammonia + Nitrate, data = meta_table[, selected_environmental_data, drop = F])

| **Variables** | **Degrees of Freedom** | **Sum of Squares** | **F** | **Pr (>F)** | **Significance** |
| --- | --- | --- | --- | --- | --- |
| pH | 1 | 0.07373 | 1.1766 | 0.2907 |  |
| Conductivity | 1 | 0.18276 | 2.9167 | 0.0184 | * |
| Chlorine | 1 | 0.14964 | 2.388 | 0.0412 | * |
| Temperature | 1 | 0.08612 | 1.3743 | 0.2053 |  |
| Dissolved oxygen | 1 | 0.11357 | 1.8124 | 0.1026 |  |
| Phosphate | 1 | 0.02585 | 0.4126 | 0.8905 |  |
| Temperature | 1 | 0.06202 | 0.9897 | 0.3816 |  |
| Ammonia | 1 | 0.07035 | 1.1227 | 0.3101 |  |
| Nitrate | 1 | 0.08086 | 1.2904 | 0.23 |  |
| Residual | 28 | 1.75452 |  |  |  |

Signif. codes: 0 ‘***’ 0.001 ‘**’ 0.01 ‘*’ 0.05 ‘.’ 0.1 ‘ ’ 1

**Table S9:** Variance Partition analyses using water chemistry/environmental parameters identified as significant being significantly associated with KO Bray-Curtis distance matrix by dbRDA analyses.

Command used:

KO_var<-varpart(KO.dist, ~Chlorine, ~Conductivity, data=metadata)

No. of explanatory tables: 2
Total variation (SS): 3.2891
No. of observations: 38

X1 = Chlorine, X2 = Conductivity

| **Variables and their combinations** | **Degrees of freedom** | **R square** | **Adjusted R square** | **Testable** |
| --- | --- | --- | --- | --- |
| [a+b] = X1 | 1 | 0.11454 | 0.08994 | TRUE |
| [b+c] = X2 | 1 | 0.11503 | 0.09044 | TRUE |
| [a+b+c] = X1+X2 | 2 | 0.20149 | 0.15586 | TRUE |
| Individual fractions |  |  |  |  |
| [a] = X1\|X2 | 1 |  | 0.06542 | TRUE |
| [b] | 0 |  | 0.02452 | TRUE |
| [c] = X2\|X1 | 1 |  | 0.06592 | TRUE |
| [d] = Residuals |  |  | 0.84414 | FALSE |

**Table S10:** Read per kilobase million (RPKM) values of all annotated KO's detected in each sample. (see excel spreadsheet).

**Table S11:** Read count per sample for KEGG modules used for DESeq2 analyses. Only modules with greater than 50% completeness or less than one block missing were. The median read count of KO's within each module was used as the module read count for DESeq2 analyses. (see excel spreadsheet).

**Table S12**: Summary of modules that were significantly higher abundance in disinfected systems as compared to non-disinfected systems.

| **Module** | **baseMean** | **log2 FoldChange** | **lfcSE** | **stat** | **pvalue** | **padj** | **Module description** | **Subcategory** | **Category** |
| --- | --- | --- | --- | --- | --- | --- | --- | --- | --- |
| M00136 | 234.06 | -27.00 | 2.95 | -9.16 | 5.03E-20 | 7.36E-19 | GABA biosynthesis, prokaryotes, putrescine => GABA | Polyamine biosynthesis | Nucleotide and amino acid metabolism |
| M00548 | 209.34 | -26.85 | 2.95 | -9.11 | 8.00E-20 | 1.09E-18 | Benzene degradation, benzene => catechol | Aromatics degradation | Secondary metabolism |
| M00367 | 202.26 | -26.54 | 2.95 | -9.01 | 2.15E-19 | 2.75E-18 | C10-C20 isoprenoid biosynthesis, non-plant eukaryotes | Terpenoid backbone biosynthesis | Carbohydrate and lipid metabolism |
| M00539 | 135.17 | -26.26 | 2.95 | -8.91 | 5.11E-19 | 5.82E-18 | Cumate degradation, p-cumate => 2-oxopent-4-enoate + 2-methylpropanoate | Aromatics degradation | Secondary metabolism |
| M00551 | 146.57 | -25.63 | 2.37 | -10.82 | 2.90E-27 | 5.40E-26 | Benzoate degradation, benzoate => catechol / methylbenzoate => methylcatechol | Aromatics degradation | Secondary metabolism |
| M00861 | 4.30 | -21.58 | 2.95 | -7.32 | 2.57E-13 | 2.03E-12 | beta-Oxidation, peroxisome, VLCFA | Fatty acid metabolism | Carbohydrate and lipid metabolism |
| M00107 | 3339.85 | -5.93 | 1.35 | -4.39 | 1.14E-05 | 6.32E-05 | Steroid hormone biosynthesis, cholesterol => prognenolone => progesterone | Sterol biosynthesis | Carbohydrate and lipid metabolism |
| M00545 | 7618.51 | -4.83 | 1.27 | -3.80 | 1.43E-04 | 6.66E-04 | Trans-cinnamate degradation, trans-cinnamate => acetyl-CoA | Aromatic amino acid metabolism | Nucleotide and amino acid metabolism |
| M00597 | 7007.83 | -3.87 | 1.04 | -3.71 | 2.04E-04 | 9.07E-04 | Anoxygenic photosystem II | Photosynthesis | Energy metabolism |
| M00568 | 4616.59 | -3.83 | 0.91 | -4.23 | 2.35E-05 | 1.27E-04 | Catechol ortho-cleavage, catechol => 3-oxoadipate | Aromatics degradation | Secondary metabolism |
| M00027 | 7825.16 | -3.73 | 1.07 | -3.50 | 4.73E-04 | 1.98E-03 | GABA (gamma-Aminobutyrate) shunt | Other amino acid metabolism | Nucleotide and amino acid metabolism |
| M00061 | 13554.83 | -3.25 | 0.82 | -3.96 | 7.57E-05 | 3.78E-04 | D-Glucuronate degradation | Other carbohydrate metabolism | Carbohydrate and lipid metabolism |
| M00631 | 12841.60 | -2.40 | 0.75 | -3.21 | 1.34E-03 | 4.82E-03 | D-Galacturonate degradation (bacteria) | Other carbohydrate metabolism | Carbohydrate and lipid metabolism |
| M00066 | 15629.00 | -2.09 | 0.54 | -3.88 | 1.03E-04 | 5.03E-04 | Lactosylceramide biosynthesis | Lipid metabolism | Carbohydrate and lipid metabolism |
| M00741 | 25451.60 | -1.98 | 0.60 | -3.31 | 9.29E-04 | 3.73E-03 | Propanoyl-CoA metabolism, propanoyl-CoA => succinyl-CoA | Other carbohydrate metabolism | Carbohydrate and lipid metabolism |
| M00307 | 27139.54 | -1.69 | 0.42 | -4.06 | 4.88E-05 | 2.56E-04 | Pyruvate oxidation, pyruvate => acetyl-CoA | Central carbohydrate metabolism | Carbohydrate and lipid metabolism |
| M00036 | 24198.24 | -1.62 | 0.40 | -4.03 | 5.59E-05 | 2.86E-04 | Leucine degradation, leucine => acetoacetate + acetyl-CoA | Branched-chain amino acid metabolism | Nucleotide and amino acid metabolism |
| M00088 | 35331.02 | -1.60 | 0.30 | -5.32 | 1.04E-07 | 6.87E-07 | Ketone body biosynthesis, acetyl-CoA => acetoacetate/3-hydroxybutyrate/acetone | Fatty acid metabolism | Carbohydrate and lipid metabolism |
| M00011 | 33315.58 | -1.12 | 0.14 | -8.00 | 1.26E-15 | 1.17E-14 | Citrate cycle, second carbon oxidation, 2-oxoglutarate => oxaloacetate | Central carbohydrate metabolism | Carbohydrate and lipid metabolism |
| M00009 | 33466.36 | -0.86 | 0.15 | -5.57 | 2.52E-08 | 1.78E-07 | Citrate cycle (TCA cycle, Krebs cycle) | Central carbohydrate metabolism | Carbohydrate and lipid metabolism |
| M00173 | 25002.47 | -0.83 | 0.23 | -3.66 | 2.52E-04 | 1.10E-03 | Reductive citrate cycle (Arnon-Buchanan cycle) | Carbon fixation | Energy metabolism |
| M00121 | 46144.04 | -0.57 | 0.11 | -5.13 | 2.93E-07 | 1.88E-06 | Heme biosynthesis, glutamate => heme | Cofactor and vitamin biosynthesis | Nucleotide and amino acid metabolism |

**Table S13**: Summary of modules that were significantly higher abundance in non-disinfected systems as compared to disinfected systems.

| **Module** | **baseMean** | **log2 FoldChange** | **lfcSE** | **stat** | **pvalue** | **padj** | **Module description** | **Subcategory** | **Category** |
| --- | --- | --- | --- | --- | --- | --- | --- | --- | --- |
| M00761 | 1255.12 | 30.00 | 1.46 | 20.55 | 7.30E-94 | 7.48E-92 | Undecaprenylphosphate alpha-L-Ara4N biosynthesis, UDP-GlcA => undecaprenyl phosphate alpha-L-Ara4N | Other carbohydrate metabolism | Carbohydrate and lipid metabolism |
| M00159 | 885.38 | 30.00 | 1.45 | 20.71 | 2.92E-95 | 5.98E-93 | V-type ATPase, prokaryotes | ATP synthesis | Energy metabolism |
| M00377 | 475.90 | 30.00 | 2.19 | 13.67 | 1.51E-42 | 4.44E-41 | Reductive acetyl-CoA pathway (Wood-Ljungdahl pathway) | Carbon fixation | Energy metabolism |
| M00620 | 786.19 | 30.00 | 2.11 | 14.22 | 7.20E-46 | 2.46E-44 | Incomplete reductive citrate cycle, acetyl-CoA => oxoglutarate | Carbon fixation | Energy metabolism |
| M00175 | 4374.13 | 30.00 | 2.07 | 14.47 | 1.79E-47 | 7.34E-46 | Nitrogen fixation, nitrogen => ammonia | Nitrogen metabolism | Energy metabolism |
| M00847 | 164.40 | 29.94 | 2.21 | 13.55 | 7.99E-42 | 2.05E-40 | Heme biosynthesis, archaea, siroheme => heme | Cofactor and vitamin biosynthesis | Nucleotide and amino acid metabolism |
| M00422 | 152.52 | 29.84 | 2.24 | 13.33 | 1.47E-40 | 3.36E-39 | Acetyl-CoA pathway, CO2 => acetyl-CoA | Methane metabolism | Energy metabolism |
| M00042 | 57.83 | 28.48 | 2.94 | 9.69 | 3.28E-22 | 5.60E-21 | Catecholamine biosynthesis, tyrosine => dopamine => noradrenaline => adrenaline | Aromatic amino acid metabolism | Nucleotide and amino acid metabolism |
| M00094 | 59.01 | 27.78 | 2.94 | 9.45 | 3.34E-21 | 5.27E-20 | Ceramide biosynthesis | Lipid metabolism | Carbohydrate and lipid metabolism |
| M00836 | 23.76 | 27.05 | 2.39 | 11.34 | 8.62E-30 | 1.77E-28 | Coenzyme F430 biosynthesis, sirohydrochlorin => coenzyme F430 | Cofactor and vitamin biosynthesis | Nucleotide and amino acid metabolism |
| M00099 | 12.71 | 26.36 | 2.94 | 8.96 | 3.24E-19 | 3.90E-18 | Sphingosine biosynthesis | Lipid metabolism | Carbohydrate and lipid metabolism |
| M00801 | 9.91 | 25.99 | 2.94 | 8.83 | 1.03E-18 | 1.07E-17 | dTDP-L-olivose biosynthesis | Other carbohydrate metabolism | Carbohydrate and lipid metabolism |
| M00803 | 9.89 | 25.99 | 2.94 | 8.83 | 1.05E-18 | 1.07E-17 | dTDP-D-angolosamine biosynthesis | Other carbohydrate metabolism | Carbohydrate and lipid metabolism |
| M00047 | 6.09 | 25.32 | 2.95 | 8.59 | 8.46E-18 | 8.26E-17 | Creatine pathway | Other amino acid metabolism | Nucleotide and amino acid metabolism |
| M00179 | 29657.67 | 19.39 | 0.98 | 19.69 | 2.82E-86 | 1.92E-84 | Ribosome, archaea | Ribosome | Genetic information processing |
| M00840 | 20672.63 | 18.87 | 0.98 | 19.28 | 8.27E-83 | 4.24E-81 | Tetrahydrofolate biosynthesis, mediated by ribA and trpF, GTP => THF | Cofactor and vitamin biosynthesis | Nucleotide and amino acid metabolism |
| M00031 | 584.40 | 11.18 | 2.05 | 5.47 | 4.61E-08 | 3.15E-07 | Lysine biosynthesis, mediated by LysW, 2-aminoadipate => lysine | Lysine metabolism | Nucleotide and amino acid metabolism |
| M00100 | 149.70 | 6.09 | 1.86 | 3.27 | 1.07E-03 | 4.22E-03 | Sphingosine degradation | Lipid metabolism | Carbohydrate and lipid metabolism |
| M00623 | 5079.42 | 5.50 | 1.51 | 3.65 | 2.67E-04 | 1.14E-03 | Phthalate degradation, phthalate => protocatechuate | Aromatics degradation | Secondary metabolism |
| M00183 | 107701.24 | 2.15 | 0.28 | 7.59 | 3.11E-14 | 2.65E-13 | RNA polymerase, bacteria | RNA polymerase | Genetic information processing |
| M00308 | 95129.26 | 2.01 | 0.33 | 6.09 | 1.10E-09 | 8.39E-09 | Semi-phosphorylative Entner-Doudoroff pathway, gluconate => glycerate-3P | Central carbohydrate metabolism | Carbohydrate and lipid metabolism |
| M00178 | 105678.14 | 1.92 | 0.26 | 7.33 | 2.27E-13 | 1.86E-12 | Ribosome, bacteria | Ribosome | Genetic information processing |
| M00157 | 98147.17 | 1.75 | 0.29 | 6.00 | 1.94E-09 | 1.42E-08 | F-type ATPase, prokaryotes and chloroplasts | ATP synthesis | Energy metabolism |
| M00855 | 20526.10 | 1.60 | 0.50 | 3.21 | 1.33E-03 | 4.82E-03 | Glycogen degradation, glycogen => glucose-6P | Other carbohydrate metabolism | Carbohydrate and lipid metabolism |
| M00141 | 91291.55 | 1.56 | 0.20 | 7.97 | 1.55E-15 | 1.38E-14 | C1-unit interconversion, eukaryotes | Cofactor and vitamin biosynthesis | Nucleotide and amino acid metabolism |
| M00053 | 75805.14 | 1.02 | 0.21 | 4.81 | 1.47E-06 | 8.62E-06 | Pyrimidine deoxyribonuleotide biosynthesis, CDP/CTP => dCDP/dCTP,dTDP/dTTP | Pyrimidine metabolism | Nucleotide and amino acid metabolism |
| M00007 | 42251.43 | 0.95 | 0.19 | 4.99 | 6.10E-07 | 3.79E-06 | Pentose phosphate pathway, non-oxidative phase, fructose 6P => ribose 5P | Central carbohydrate metabolism | Carbohydrate and lipid metabolism |
| M00048 | 52586.77 | 0.87 | 0.19 | 4.49 | 7.23E-06 | 4.12E-05 | Inosine monophosphate biosynthesis, PRPP + glutamine => IMP | Purine metabolism | Nucleotide and amino acid metabolism |
| M00052 | 85948.16 | 0.83 | 0.22 | 3.84 | 1.21E-04 | 5.78E-04 | Pyrimidine ribonucleotide biosynthesis, UMP => UDP/UTP,CDP/CTP | Pyrimidine metabolism | Nucleotide and amino acid metabolism |
| M00168 | 44967.05 | 0.73 | 0.23 | 3.25 | 1.17E-03 | 4.53E-03 | CAM (Crassulacean acid metabolism), dark | Carbon fixation | Energy metabolism |
| M00035 | 52458.61 | 0.71 | 0.21 | 3.32 | 8.87E-04 | 3.64E-03 | Methionine degradation | Cysteine and methionine metabolism | Nucleotide and amino acid metabolism |
| M00002 | 50244.03 | 0.68 | 0.14 | 4.91 | 9.09E-07 | 5.48E-06 | Glycolysis, core module involving three-carbon compounds | Central carbohydrate metabolism | Carbohydrate and lipid metabolism |
| M00125 | 46145.15 | 0.54 | 0.14 | 3.75 | 1.76E-04 | 8.04E-04 | Riboflavin biosynthesis, GTP => riboflavin/FMN/FAD | Cofactor and vitamin biosynthesis | Nucleotide and amino acid metabolism |
| M00049 | 63394.75 | 0.49 | 0.15 | 3.23 | 1.22E-03 | 4.62E-03 | Adenine ribonucleotide biosynthesis, IMP => ADP,ATP | Purine metabolism | Nucleotide and amino acid metabolism |
| M00050 | 61091.85 | 0.45 | 0.14 | 3.22 | 1.30E-03 | 4.82E-03 | Guanine ribonucleotide biosynthesis IMP => GDP,GTP | Purine metabolism | Nucleotide and amino acid metabolism |

**Table S14:** Summary statistics for metagenome assembled genomes (MAGs) extracted from the metagenome assemblies. Metagenome assembled genomes (MAG) were finalized after dereplication using dRep (https://github.com/MrOlm/drep) using all MAGs assembled in this study. As a result, the name assigned to a MAG does not represent sampling location it was assembled from. MAGs were assigned taxonomy using the Genome taxonomy database (GTDB-TK: https://gtdb.ecogenomic.org/) version 0.1.3. The completeness and redundancy of MAGs was estimated using CheckM (https://github.com/Ecogenomics/CheckM/wiki) version 1.0.7. Only MAGs >50% completeness and <10% redundancy were included in the study. The genome statistics were estimated using Prokka. The coding density was estimated by dividing the cumulative length of coding sequences (CDS) divided by the length of the MAG. The MAGs were assigned four categories, "D-only", "ND-only", "Both", and "Other". "D-only" was assigned to MAGs detected in >20% of disinfected samples and not detected in non-disinfected samples. "ND-only" was assigned to MAGs detected in <20% of disinfected samples and not detected of non-disinfected samples. "Both" was assigned to MAGs that were detected in >20% of disinfected and non-disinfected samples. "Other" was assigned to MAGs that did not fall in either of the above three classes. (see excel spreadsheet).

**Table S15:** Metagenome assembled genomes were considered detected in a sample if 25% of the nucleotides were covered by reads mapped from the sample. Reads were mapped using bwa mem (v 0.7.3a) and the coverage per base was determined using bedtools genomcov. "Proportion of genome detected sheet" indicates what proportion of the nucleotides in a MAG has atleast 1x coverage "MAG_detect_nondetect" shows bins that were considered detected (shown by "1") and non-detected (shown by "0") in each sample.

**Table S16:** KEGG modules and their completeness estimates within each MAG assembled as part of this study.

**Table S17**: KOfamScan based annotation of MAGs indicating presence/absence of detected KEGG orgthologies (KO) in each MAG.
